# Transcriptional dynamics of sleep deprivation and subsequent recovery sleep in the male mouse cortex

**DOI:** 10.1101/2024.08.20.607983

**Authors:** Alexander Popescu, Caitlin Ottaway, Kaitlyn Ford, Taylor Wintler Patterson, Ashley Ingiosi, Elizabeth Medina, Stephanie C. Hicks, Kristan Singletary, Lucia Peixoto

**Author notes:** Correspondence: Lucia Peixoto. Denotes authors contributed equally (Alexander Popescu, Caitlin Ottaway).

## Abstract

Sleep is an essential, tightly regulated biological function. Sleep is also a homeostatic process, with the need to sleep increasing as a function of being awake. Acute sleep deprivation (SD) increases sleep need, and subsequent recovery sleep (RS) discharges it. SD is known to alter brain gene expression in rodents, but it remains unclear which changes are linked to sleep homeostasis, SD-related impairments, or non-sleep-specific effects. To investigate this question, we analyzed RNA-seq data from adult wild-type male mice subjected to 3 and 5-6 hours of SD and 2 and 6 hours of RS after SD. We hypothesized molecular changes associated with sleep homeostasis mirror sleep pressure dynamics as defined by brain electrical activity, peaking at 5-6 hours of SD, and are no longer differentially expressed after 2 hours of RS. We report 5-6 hours of SD produces the largest effect on gene expression, affecting approximately half of the cortical transcriptome, with most differentially expressed genes (DEGs) downregulated. The majority of DEGs normalize after 2 hours of RS and are involved in redox metabolism, chromatin regulation, and DNA damage/repair. Additionally, RS affects gene expression related to mitochondrial metabolism and Wnt-signaling, potentially contributing to its restorative effects. DEGs associated with cholesterol metabolism and stress response do not normalize within 6 hours and may be non-sleep-specific. Finally, DEGs involved in insulin signaling, MAPK signaling, and RNA-binding may mediate the impairing effects of SD. Overall, our results offer insight into the molecular mechanisms underlying sleep homeostasis and the broader effects of SD.

**New & Noteworthy:** This study investigates different time points of sleep deprivation and recovery sleep to better understand the molecular processes influenced by sleep and lack of sleep. This study highlights redox metabolism, chromatin regulation, and DNA damage/repair as molecular mechanisms linked to sleep homeostasis while showing the effects of stress are probably non-sleep-specific based on transcriptional dynamics.

## Introduction

Sleep is a tightly regulated essential biological function. Sleep regulation is controlled by two processes, a circadian process that determines the timing of sleep and a homeostatic process that determines sleep need as a function of being awake (sleep pressure) (1). Acute sleep deprivation (SD) increases sleep pressure, and subsequent sleep, termed recovery sleep (RS), discharges it. The dynamics of this are physiologically well understood based on measures of brain electrical activity and known to be under genetic control (2). However, the cellular and molecular basis of homeostatic sleep pressure remains poorly understood.

Sleep and sleep loss have consistently been shown to regulate gene expression throughout the brain in rodents (3–14). SD impacts the frontal cortex transcriptome (13), with studies identifying specific gene-associated cellular localizations (10) and functions (8, 13) affected by SD. However, which proportion of those changes are directly linked to increased sleep pressure remains underexplored. In particular, there has been little work on transcriptional dynamics across both short and prolonged amounts of RS. Additionally, SD impairs various cognitive functions, including working memory (15) and spatial learning (16), yet the dynamics of recovery for genes associated with these functions are underexplored. We hypothesize that transcriptional changes after SD, which form the basis of sleep homeostasis, respond dynamically to subsequent RS with similar dynamics as brain activity measures of sleep pressure. Our lab previously studied these dynamics utilizing microarrays, finding that the effects of acute SD are quickly reversible (8). These findings align biologically with the primary discharge of sleep pressure in the first 2 hours of RS. However, the molecular basis of sleep homeostasis remains poorly described. As such, exploring the dynamics of SD and RS is vital to further understanding the molecular basis of the accumulation of sleep pressure.

To address this question, we integrated data from our lab with a publicly available RNA-seq dataset to investigate transcriptional dynamics at the gene level across multiple SD and RS time points in the cortex of adult wild-type male mice. We report a largely repressive effect of SD that affects approximately half of the cortical transcriptome as we have previously reported (13), which is maximal at 5-6 hours when sleep pressure is highest. This study identifies redox metabolism, chromatin regulation, and DNA damage/repair as likely molecular mechanisms of sleep homeostasis. Finally, we provide potential insight into the functions that may mediate the impairments seen during SD, as well as non-sleep-specific effects of cellular stress on the transcriptional response to SD.

## Materials and Methods

### Experimental design and animals

Most of the data in this study were extracted from two previously published RNA-seq datasets: GSE140345 (14) and GSE113754 (17). For detailed methodology regarding these studies, please see the respective citation. Described here is the protocol for mice subjected to 5 hours of sleep deprivation (SD5) followed by 2 hours of subsequent recovery sleep (RS2) and their corresponding home cage controls (HC7). All experimental procedures were approved by the Institutional Animal Care and Use Committee of Washington State University and conducted in accordance with National Research Council guidelines and regulations for experiments in live animals.

Adult male wild-type (WT) C57BL/6J mice (8–12 weeks old) were used in all studies to allow for comparison with the publicly available datasets. SD was performed via gentle handling as previously described (17). Briefly, mice were individually housed in standard cages at 24 ± 1°C on a 12:12 hour light/dark cycle with food and water *ad libitum* seven days before tissue collection. Mice were assigned to either SD with subsequent RS (SD + RS, n = 5 independent animals) or home cage (HC, n = 5) control conditions. Beginning at light onset, zeitgeber time 0 (ZT0), the mice in the SD + RS group were kept awake for 5 hours and then allowed to sleep for 2 hours. Mice in the HC group served as circadian controls and were left undisturbed in their HC for 7 hours from ZT0 to ZT7.

### Tissue collection

Without prior anesthesia, mice were immediately sacrificed at ZT7 by cervical dislocation and decapitation, alternating between the SD + RS and HC groups. Frontal cortex tissue was dissected on a cold block, flash-frozen in liquid nitrogen, and stored at –80°C until processing (11, 17). This protocol was repeated over five days, with one sample from each treatment group collected daily and all tissue collected within the first 15 minutes of the hour.

### RNA isolation and sequencing

Tissue was homogenized in Qiazol buffer (Qiagen, Hilden, Germany) using a TissueLyser (Qiagen), and RNA was extracted using the Qiagen RNeasy kit. The integrity of total RNA was verified using Fragment Analyzer (Advanced Analytical Technologies, Inc, Ankeny, IA) with the High Sensitivity RNA Analysis Kit (Advanced Analytical Technologies, Inc). The TruSeq Stranded mRNA Library Prep Kit (Illumina, San Diego, CA) was used for RNA library preparation. Briefly, mRNA was isolated from 2.5 mg of total RNA using poly-T oligo attached to magnetic beads and then subjected to fragmentation, followed by cDNA synthesis, dA-tailing, adaptor ligation, and PCR enrichment. RNA library sizes were assessed by Fragment Analyzer with the High Sensitivity NGS Fragment Analysis Kit (Advanced Analytical Technologies, Inc). The concentrations of RNA libraries were measured by the StepOnePlus Real-Time PCR System (ThermoFisher Scientific, San Jose, CA) with the KAPA Library Quantification Kit (Kapa Biosystems, Wilmington, MA). The libraries were diluted to 2 nM with Tris buffer (10 mM Tris-HCl, pH 8.5) and denatured with 0.1 M NaOH. Eighteen pM libraries were clustered in a high-output flow cell using the HiSeq Cluster Kit v4 on a cBot (Illumina). After cluster generation, the flow cell was loaded onto the HiSeq 2500 for sequencing using the HiSeq SBS kit v4 (Illumina). DNA was paired-end sequenced with a read length of 100 bp and an average sequencing depth of 54 million read pairs per sample. HiSeq Control Software (v. 2.2.68) was used for base calling, and raw bcl files were converted to FASTQ files using bcl2fastq (v. 2.17.1.14). Adapters were trimmed from the FASTQ files during the conversion. All library preparation and sequencing steps were performed by the WSU Spokane Genomics Core.

### Data integration

The RS2 and HC7 time points were integrated with samples from GSE140345 and GSE113754 (**Figure 1**). Altogether, the following time points were included: 3, 5, and 6 hours of SD (SD3, SD5, SD6); 5 hours of SD followed by 2 hours of RS (RS2); and 6 hours of SD followed by 6 hours of RS (RS6). For each time point, there were time-matched home cage controls (HC3, HC5, HC6, HC7, HC12) to account for variations in circadian timing. Samples for the SD5 and HC5 time points were generated in GSE113754 and extracted from frontal cortex tissue. Samples for the SD3, HC3, SD6, HC6, RS6, and HC12 time points were generated in GSE140345 and extracted from cerebral cortex tissue. There were n = 3 independent animals for the SD3, HC3, SD6, and RS6 time points and n = 5 independent animals for the SD5, HC5, HC6, RS2, HC7, and HC12 time points.

**Figure 1.**
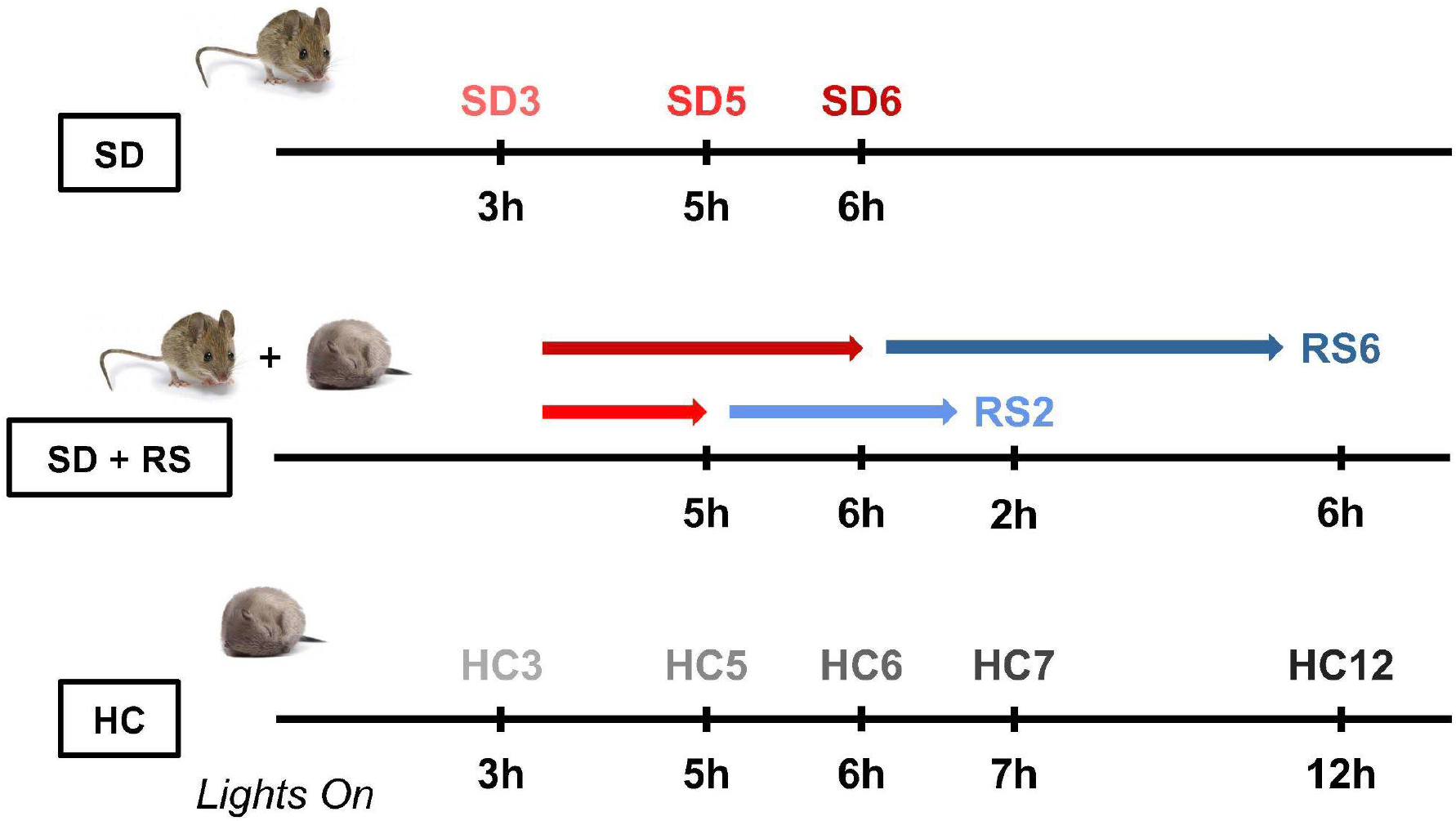
Summary of experimental time points in this study. Top: SD3 (n = 3), SD5 (n = 5), SD6 (n = 3); 3, 5, and 6 hours of sleep deprivation (SD). Middle: RS2 (n = 5), RS6 (n = 3); 2 and 6 hours of recovery sleep (RS) after SD5 and SD6, respectively. Bottom: HC3 (n = 3), HC5 (n = 5), HC6 (n = 5), HC7 (n = 5), HC12 (n = 5); circadian home cage (HC) controls for each SD and RS time point. Red colors indicate SD, blue colors indicate RS, and gray colors indicate HC.

### Transcript quantification

Raw sequencing reads were quantified using Salmon (v. 1.8.0) (18). Briefly, the reference genome and transcriptome annotation files were downloaded from GENCODE (release M28, genome assembly GRCm39). The genome targets, which served as decoys, and the concatenated list of transcriptome targets were used to build the index with the following parameters: `--gencode` and `-k 31`. For the quantification step, the following parameters were used: `--libType A` and `--numBootstraps 30`. The Tximeta R/Bioconductor package (v. 1.18.0) (19) was used to import transcript quantification data from Salmon and summarize transcript-level quantifications to the gene level. Other packages from the R/Bioconductor project (v. 3.17.1) (20, 21) were used throughout the data analysis pipeline.

### Differential expression analysis

To account for inflated log2 fold change (log2FC) values associated with lowly expressed genes, the matrix of gene counts was filtered to remove genes with less than ten reads across more than five samples. Selecting five samples made sense for our dataset as the majority of experimental conditions had five replicates. Upper-quartile between-lane normalization was implemented using the EDASeq package (v. 2.34.0) (22) to correct for library size. Unwanted variation was estimated using the RUVSeq package (v. 1.34.0) (23). k = 3 was needed to adequately correct for batch effects. A set of 3422 genes (Additional File 4 from (8), adj. p-value > 0.9) less likely to be affected by SD were used as negative controls for RUVs. These genes were obtained from an independent microarray study after 5–6 hours of SD (8) and will be referred to as *empirical* negative controls hereafter.

Differential gene expression analysis with a mixed effects model (QLF plus unwanted factors estimated during RUVs normalization) was performed using the edgeR package (v. 3.38.1) (24) with FDR < 0.05. The SD5 and SD6 time points and the HC5 and HC6 time points were grouped to obtain a single estimate of the effects of a long (5–6 hours) SD duration and to remove the batch effects due to different laboratories and brain regions (cortex versus frontal cortex). Given that sleep pressure accumulation plateaus between 5-6 hours of SD (2), this approach likely captures gene expression changes at maximum sleep pressure after SD. The following pairwise comparisons were selected, with differential expression being determined relative to HC controls: SD3 vs. HC3, SD5–6 vs. HC5–6, RS2 vs. HC7, and RS6 vs. HC12.

A set of 670 positive control genes (Additional File 2 from (8), adj. p-value < 0.01) assembled independent of lab, technology, and brain region following 5–6 hours of SD were used to assess the reproducibility of results from differential expression. Approximately 200 genes (Additional File 4 from (8), adj. p-value < 0.01) previously reported to be differentially expressed at 2 or 6 hours of RS following 5–6 hours of SD in the prefrontal cortex were also used. As with the empirical negative controls, probe IDs were mapped to Ensembl IDs, which were then annotated using Ensembl release 105 (25). Probe IDs mapping to non-unique Ensembl IDs were discarded, and Ensembl IDs with no associated gene name following annotation were removed. All positive and empirical negative controls can be found in **Supplemental Table S1**.

In addition to the FDR cutoff, a log2 counts per million (log2CPM) filter was also applied to filter for differentially expressed genes. Log2CPM > 0 was chosen to optimize the recovery of SD positive controls while simultaneously minimizing the recovery of empirical negative controls. This led to over 150 fewer empirical negative controls in the dataset, with only nine positive controls lost (**Supplemental Table S2**).

### Functional enrichment analysis

The four lists of differentially expressed genes from the four pairwise comparisons were annotated with basic gene description information and separated into upregulated and downregulated lists by log2FC. UpSet plots were generated using the UpSetR (v. 1.4.0) (26) and ComplexUpset (v. 1.3.3) packages and were used to visualize all possible intersections across time points separately for upregulated and downregulated genes. Sufficiently large (at least 100 genes) and biologically relevant intersections were selected for functional enrichment analysis. Gene lists for each of the three levels of recovery regardless of the amount of SD required for differential expression (normalize within 2 hours; require 2–6 hours to normalize; require more than 6 hours to normalize) were obtained by taking the respective union (indicated with U) across SD3 and SD3/SD5–6. These lists were also selected for functional enrichment analysis.

Functional annotation of subsets of genes that fit one of the three patterns of recovery (separating by 3 or 5–6 hours of SD) or that were unique to the RS2 or RS6 time points was performed using the Database for Annotation, Visualization, and Integrated Discovery v2021 (DAVID) (27, 28) with DAVID Knowledgebase v2023q3. UniProt (https://www.uniprot.org/) biological process and molecular function terms and Kyoto Encyclopedia of Genes and Genomes (KEGG) pathways (https://www.genome.jp/kegg/pathway.html) were selected as annotation categories. An enrichment p-value (EASE score) threshold of 0.05 was used, with enrichment being determined relative to the filtered list of 18872 expressed genes. Clustering of terms was performed using initial and final group memberships of 3 and a similarity threshold of 0.20 to allow for inclusive clustering. Clustered and unclustered terms were visualized with bubble plots generated using the ggplot2 (v. 3.5.1) package. Terms within a given cluster were condensed into a single informative cluster name by taking the geometric mean of their fold enrichment values. Hub genes were defined by identifying genes that appeared in every individual term for a functional annotation cluster.

## Results

### The majority of transcriptional changes require 5–6 hours of sleep deprivation and are no longer detected after 2 hours of recovery sleep

It is known that acute SD influences genome-wide gene expression in the brain (3–14). This molecular response is, in part thought to underlie the basis of the homeostatic response to acute sleep loss. However, how different amounts of SD and subsequent recovery sleep (RS) influence the dynamics of gene expression is not well understood. Understanding how and when different genes and their associated biological functions and pathways normalize can help explain the molecular basis of sleep homeostasis. In this study, we integrated data obtained after different amounts of acute SD (3 and 5-6 hours) and subsequent RS (2 and 6 hours) from independent studies to define the ability of SD and RS to dynamically influence gene expression in the cortex of adult male mice (see Methods, **Figure 1**).

Figure 2. shows the results of normalization and differential expression analysis across all time points. The initial implementation of upper-quartile (UQ) normalization to correct for library size did not provide adequate correction for batch effects (**Supplemental Figure S1**). Specifically, the lab where samples were collected accounted for nearly half of the variation in the dataset (∼48%, PC1) (**Supplemental Figure S1A**). Thus, we used RUVseq to estimate factors of unwanted variation in our UQ-normalized dataset to better correct for confounding technical and biological factors, as in our previous work (8, 11, 29). Following *remove unwanted variation* (RUV) normalization, the main factors driving variance in the data are the amount of sleep (∼14%, PC1) and circadian time (∼6%, PC2) (**Figure 2A**). As expected, samples clustered from left to right by the experimental treatment (HC control, SD + RS, or SD only) and hours of sleep. Additionally, RUV normalization resulted in dynamic ranges that were more comparable between samples (**Figure 2B**). Following normalization, we conducted differential expression analysis. After applying a cutoff on expression level (see Methods), we identified 2315 differentially expressed genes (DEGs) after 3 hours of SD and 7493 DEGs after 5-6 hours of SD (**Figure 2C–D**). We obtained a list of positive controls genes that were previously shown to be altered after SD (8). When compared to the list of genes differentially expressed in the SD5–6 comparison, nearly 90% of positive control genes were recovered. After both 3 and 5–6 hours of SD, there were more downregulated (SD3: 1373, SD5–6: 4390) than upregulated (SD3: 942, SD5–6: 3103) DEGs. Following 2 hours of RS, 3908 genes were differentially expressed, and after 6 hours of RS, 1989 genes were differentially expressed (**Figure 2E–F**). Whereas SD had a repressive effect on transcription, RS preferentially upregulated DEGs. For the RS6 time point, there were more upregulated than downregulated DEGs (1104 versus 885). The complete list of DEGs for each time point can be found in **Supplemental Table S3**.

**Figure 2.**
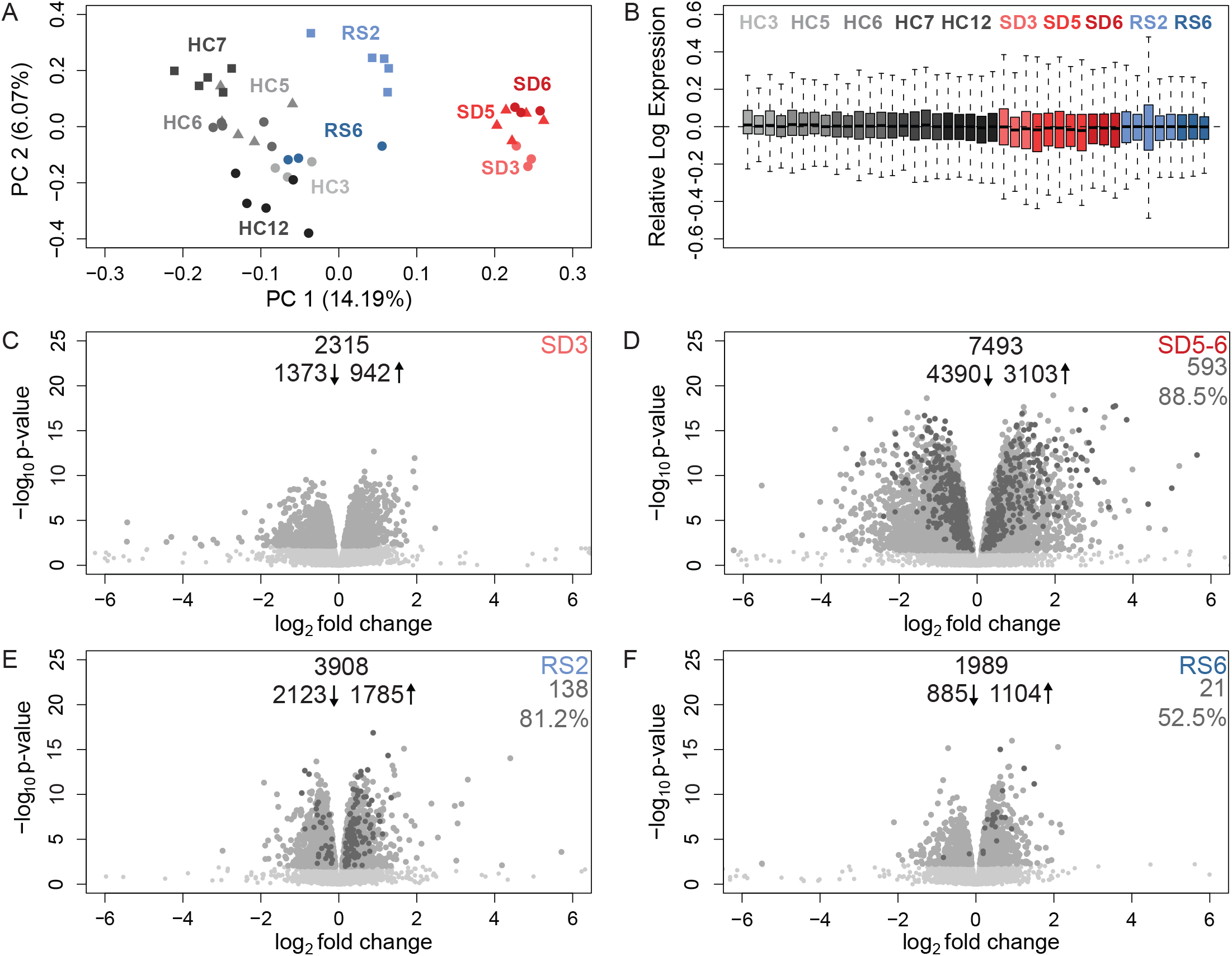
Sleep deprivation after 5-6 hours results in the largest number of differentially expressed genes compared to other time points. A) Principal component analysis following RUVs normalization with k = 3 unwanted factors. Circles represent samples from (30), triangles represent samples from (17), and squares represent new samples from our group. Home cage (HC) samples are in shades of gray, sleep deprivation (SD) samples are in shades of red, and recovery sleep (RS) samples are in shades of blue. Amount of sleep (PC1, 14.19%) and circadian time (PC2, 6.07%) account for the most variability in the dataset. B) Relative log expression for each sample and condition. Color code as in A. C– F) Volcano plots following differential expression analysis on RUVs normalized counts. C) SD3 vs HC3, D) SD5–6 vs HC5–6, E) RS2 vs HC7, F) RS6 vs HC12. Expressed genes are in light gray, differentially expressed genes (DEGs) are in gray, and positive controls are in dark gray. Numbers of downregulated DEGs upregulated DEGs, and total DEGs are reported in the top center for each comparison. The number and percentage of positive controls recovered are reported in the top right for each comparison.

To define the time point(s) at which each gene was differentially expressed in the same direction (i.e., either upregulated or downregulated) across the 4 SD and RS time points, we intersected upregulated and downregulated DEGs separately. **Figure 3** shows biologically meaningful intersections containing at least 100 DEGs (all patterns can be found in **Supplemental Figure S2**). Intersections among upregulated DEGs are shown in **Figure 3A**, and intersections among downregulated DEGs are shown in **Figure 3B**. The largest number of upregulated DEGs (see red box) were genes that returned to their baseline level of expression (i.e., normalized) within 2 hours of RS. The majority of these genes required 5-6 hours of SD (1149) to be differentially expressed, whereas the rest were already present after 3 hours of SD (407). There was a similar pattern among downregulated genes, with 2420 (1898 + 522) genes that normalized within 2 hours. As indicated by the dark magenta box, there were 966 (692 + 274) upregulated genes and 1441 (931 + 510) downregulated genes that required at least 2 hours but less than 6 hours to normalize. As with genes that normalized the fastest, most of these genes were not yet differentially expressed after only 3 hours of SD. The third pattern of recovery we considered is represented by the lavender box and included genes that need at least 6 hours of RS to no longer be differentially expressed. 338 (220 + 118) upregulated genes and 302 (185 + 117) downregulated genes fell into this category. Lastly, the dark blue box indicates genes differentially expressed with RS only and not SD. Among upregulated genes, more were unique to the RS6 time point (463) than the RS2 time point (423). There were fewer downregulated genes in this pattern (691 = 356 + 335). The complete list of DEGs for each intersection can be found in **Supplemental Table S4**.

**Figure 3.**
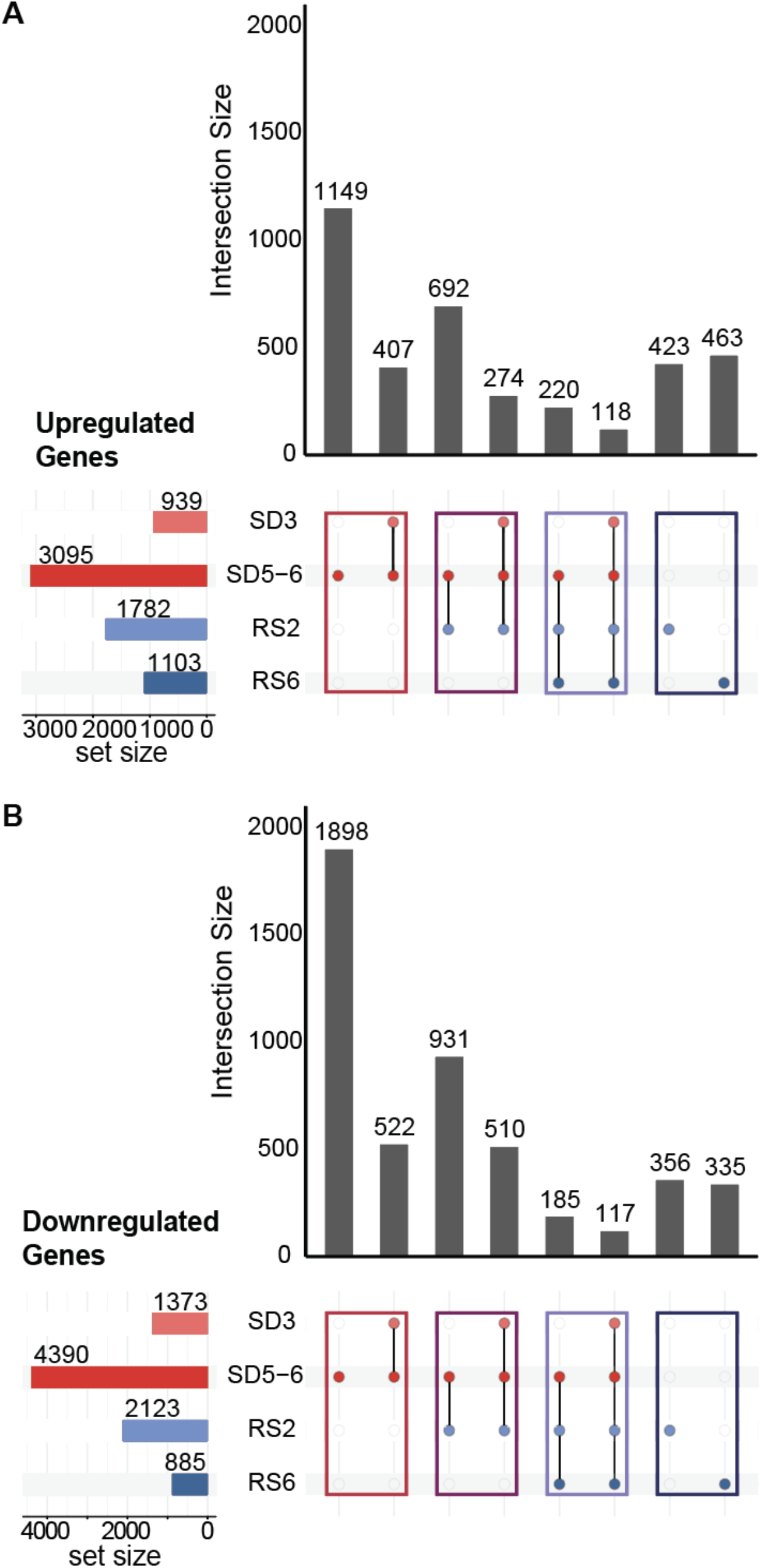
Combinatorial patterns of differential gene expression across varied amounts of sleep deprivation and recovery sleep. UpSet plots of the intersections across the SD3, SD5–6, RS2, and RS6 time points selected for functional enrichment analysis. Lists of differentially expressed genes (DEGs) were intersected separately for A) upregulated genes and B) downregulated genes. The total number of DEGs per time point is shown on the set size rows. The vertical bars represent the number of DEGs belonging to the subset depicted by the colored dots in the intersection matrix. The four colored boxes (red, dark magenta, lavender, and dark blue) indicate the four pairs of intersections shown in Figures 4–7.

Subsequently, we wanted to understand which pathways or biological functions were differentially affected by varying amounts of SD and RS. We performed a functional enrichment analysis on the intersections specified in **Figure 3** using DAVID. To better understand the functions associated with different patterns of recovery, we also performed enrichment on the union across SD3 and SD5-6 for each of the first three colored boxes (red, dark magenta, and lavender). The enrichment of Kyoto Encyclopedia of Genes and Genomes (KEGG) pathways and UniProt biological processes (BP) and molecular functions (MF) was determined relative to the matrix of expressed genes. KEGG, BP, and MF terms, herein referred to as functional terms, were clustered based on the overlap of genes. This analysis produced significantly enriched (p-value < 0.05) clusters of functional terms and unclustered terms. “Hub” genes, shown in bold in Figures 4-7, mediate clustering (see Methods). We also highlight positive control genes replicated from previous studies and representative genes from unclustered terms. The full results of functional enrichment for each intersection and union can be found in **Supplemental Table S5**.

**Figure 4.**
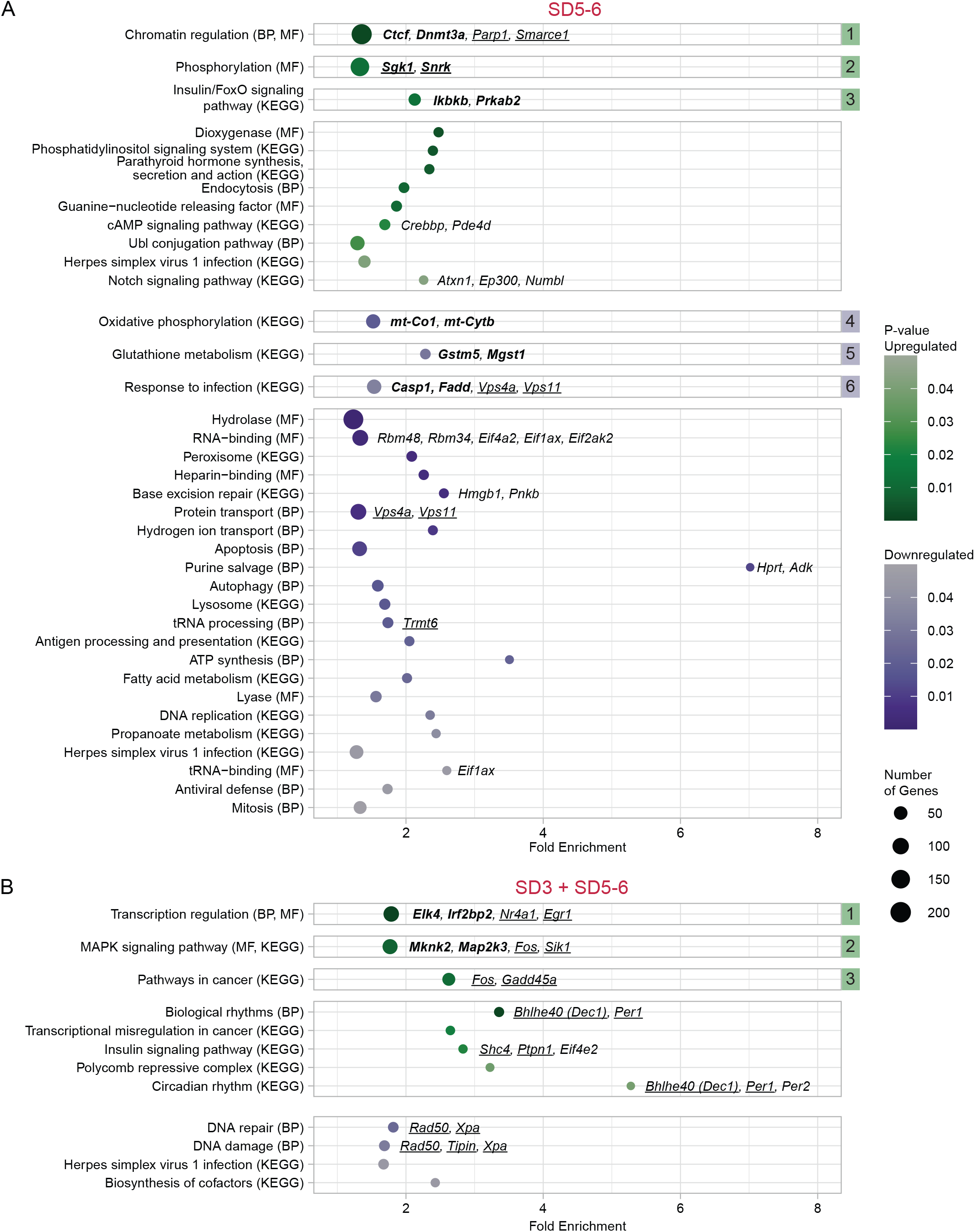
DEGs after SD that are no longer detected after 2 hours of RS are involved in chromatin regulation, oxidative phosphorylation, and DNA damage/repair. Bubble plots of the results of functional enrichment for A) genes differentially expressed only at SD5–6 and B) genes differentially expressed only at SD3 and SD5–6. Enriched (modified Fisher’s Exact p-value < 0.05) functional annotation terms or clusters are displayed vertically and plotted as circles. Clusters are denoted by numbered square boxes to the right of the figure. Functional terms are either Kyoto Encyclopedia of Genes and Genomes (KEGG) pathways, UniProt biological processes (BP), or UniProt molecular functions (MF). Circle size represents the number of genes in a term or a cluster. Upregulated functional terms are shown in green and downregulated terms in purple, with darker shades representing smaller p-values (or the geometric mean of p-values for clusters). Fold enrichment (or the geometric mean of the fold enrichment for clusters) is shown on the x-axis. Hub genes are in bold. Positive control genes are underlined. Representative genes are in regular italics.

We first examined what functional terms were enriched in genes affected by SD but no longer differentially expressed after 2 hours of RS since this group comprised the largest number of genes (**Figure 4**; red box in **Figure 3**). Terms for upregulated genes requiring 5–6 hours of SD formed clusters enriched in chromatin regulation (e.g., *Ctcf, Dnmt3a, Parp1, Smarce1*), phosphorylation (e.g., *Sgk1, Snrk*), and insulin/FoxO signaling (e.g., *Ikbkb, Prkab2*) (**Figure 4A**). Upregulated genes were also enriched in cyclic adenosine monophosphate (cAMP) signaling (e.g., *Crebbp, Pde4d*) and Notch signaling (e.g., *Atxn1, Ep300, Numbl*), which did not cluster with other functional terms. Clusters of downregulated genes after 5–6 hours of SD were enriched in oxidative phosphorylation (e.g., *mt-Co1, mt-Cytb*), glutathione metabolism (e.g., *Gtsm5, Mgst1*), and response to infection (e.g., *Casp1, Fadd, Vps4a, Vps11*). Downregulated genes also participated in RNA-binding (e.g., *Rbm48, Rbm34, Eif4a2, Eif1ax, Eif2ak2*), base excision repair (e.g., *Hmgb1, Pnkb*), protein transport (e.g., *Vps4a, Vps11*), tRNA processing (e.g., *Trmt6*), tRNA-binding (e.g., *Eif1ax*), and purine salvage (e.g., *Hprt, Adk*), which was the term with the highest fold enrichment. Notably, multiple downregulated terms and clusters, namely response to infection, hydrolase, RNA-binding, herpes simplex virus 1 infection, tRNA-binding, antiviral defense, and protein biosynthesis, as well as the upregulated cluster insulin signaling, were enriched in eukaryotic initiation factors (Eifs). Overall, chromatin regulation, insulin/FoxO signaling, and metabolism (specifically protein synthesis and energy metabolism) were the primary molecular signatures associated with transcriptional changes that require 5–6 hours of SD but normalize within 2 hours of RS.

Subsequently, we investigated functional terms enriched in genes that were differentially expressed after only 3 hours of SD, continued to be present after 5–6 hours, and normalized within 2 hours of RS (**Figure 4B**). The clusters of upregulated functional terms were transcription regulation (e.g., *Elk4, Irf2bp2, Nr4a1, Egr1*), the mitogen-activated protein kinase (MAPK) signaling pathway (e.g., *Mknk2, Map2k3, Fos, Sik1*), and pathways in cancer (e.g., *Fos, Gadd45a*). Upregulated terms that did not cluster included biological/circadian rhythms (e.g., *Bhlhe40 (Dec1), Per1*) and insulin signaling (e.g., *Shc4, Ptpn1, Eif4e2*). Circadian rhythm was the term with the highest fold enrichment. Immediate early genes (e.g., *Fos, Egr1*) and circadian genes (e.g., *Per1, Per2*) were present in multiple terms. Terms for downregulated genes did not cluster and included DNA damage/repair (e.g., *Rad50, Tipin, Xpa*). Functional enrichment analysis on the union of all genes affected by SD that normalized before 2 hours of RS revealed metabolism (e.g., *Gstp1*) and protein biosynthesis (e.g., *Eifs*) as downregulated (**Supplementary Figure S3**).

Next, we analyzed functional terms for genes affected by SD that were no longer differentially expressed after 6 hours of RS (**Figure 5**; dark magenta box in **Figure 3**). Terms for upregulated genes requiring 5–6 hours of SD formed clusters enriched in mTOR signaling (e.g., *Pik3ca, Pik3r1*) and cell adhesion (e.g., *Pik3ca, Pik3r1, Crk, Vav3*) (**Figure 5A**). Upregulated genes also belonged to circadian rhythms (e.g., *Clock, Creb1, Npas2*) and stress response (e.g., *Ahsa2, Hsf1, Hspe1*). Clusters of downregulated functional terms included cholesterol biosynthesis (e.g., *Mvd, Msmo1*), ion transport (e.g., *Kcnj4, Pllp*), cell signaling (e.g., *Kcnj4, Gnai2, Prkaca*), and glucose metabolism (e.g., *Ugp2*). Downregulated terms that did not cluster included glycogen biosynthesis (e.g., *Agl, Akt1, Gbe1*). For genes already differentially expressed after 3 hours of SD, clusters of upregulated functional terms included RNA-binding (e.g., *Srsf7, Tra2a, Rbm12b1, Ythdc1*) and stress response (e.g., *Dnajc3, Hspa1b, Hspa5*) (**Figure 5B**). Some upregulated terms, like chromatin regulator (e.g., *Chd9*), did not cluster.

**Figure 5.**
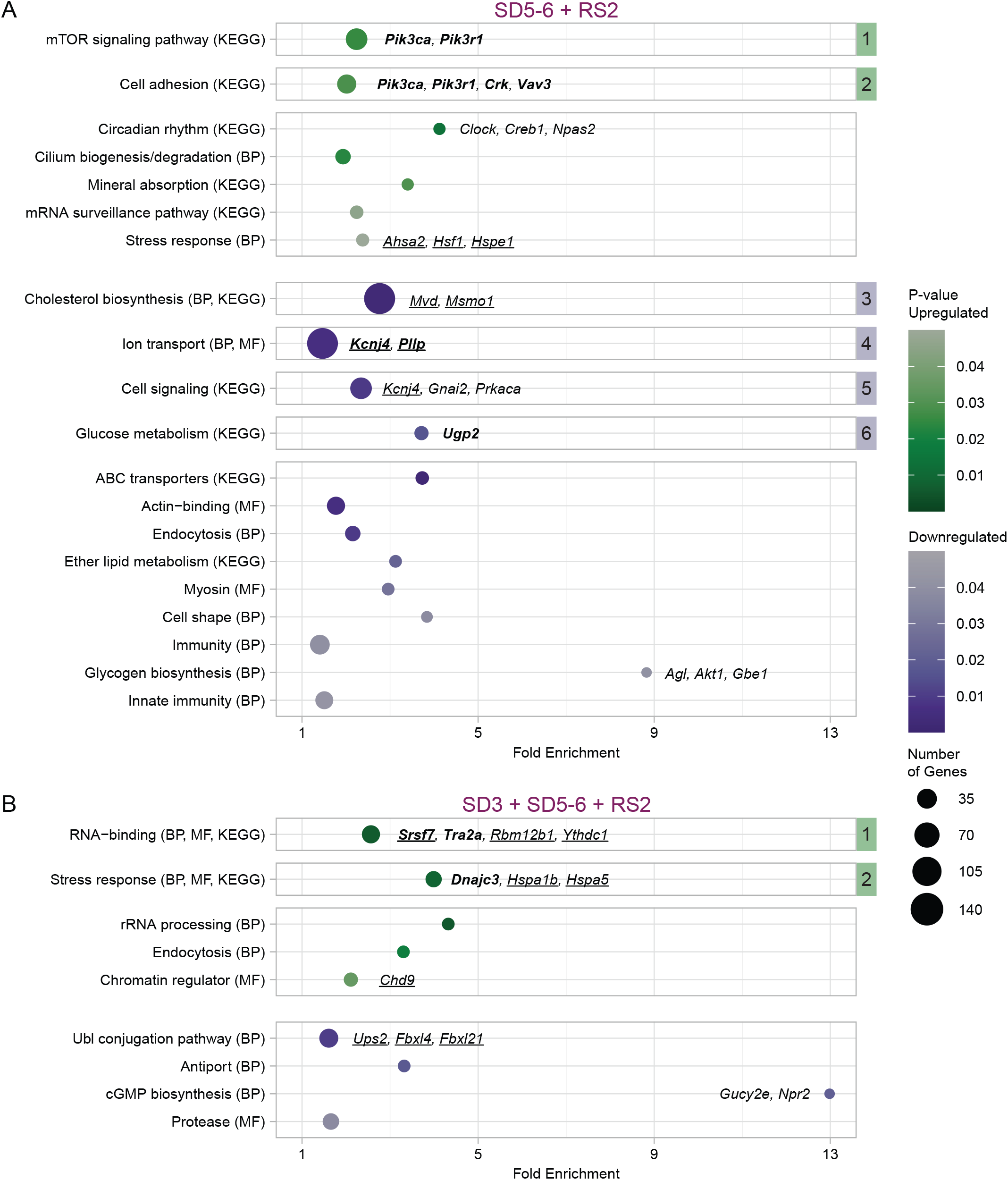
DEGs after SD that are no longer detected after 2–6 hours of RS are involved in cholesterol biosynthesis, mTOR signaling, stress response, and ion transport. Bubble plots of the results of functional enrichment for A) genes differentially expressed only at SD5–6 and RS2 and B) genes differentially expressed at SD3, SD5–6, and RS2, but not RS6. Enriched (modified Fisher’s Exact p-value < 0.05) functional annotation terms or clusters are displayed vertically and plotted as circles. Clusters are denoted by numbered square boxes to the right of the figure. Functional terms are either Kyoto Encyclopedia of Genes and Genomes (KEGG) pathways, UniProt biological processes (BP), or UniProt molecular functions (MF). Circle size represents the number of genes in a term or a cluster. Upregulated functional terms are shown in green and downregulated terms in purple, with darker shades representing smaller p-values (or the geometric mean of p-values for clusters). Fold enrichment (or the geometric mean of the fold enrichment for clusters) is shown on the x-axis. Hub genes are in bold. Positive control genes are underlined. Representative genes are in regular italics.

Downregulated functional terms were all unclustered and included ubiquitin-like (Ubl) protein conjugation (e.g., *Usp2, Fbxl4, Fbxl21*) and cGMP biosynthesis (e.g., *Npr2, Gucy2e*), which was the term with the highest fold enrichment. Functional enrichment analysis on the union of all genes affected by SD that normalized within 6 hours highlighted stress response (e.g., *Dnajc3, Hspa2b*) and cholesterol biosynthesis (e.g., *Mvd*) as important molecular signatures (**Supplementary Figure S4**).

### Transcriptional changes after SD that are no longer detected after 6 or more hours of RS are involved in stress response and metabolism

Next, we analyzed functional terms for genes affected by SD that were no longer differentially expressed after 6 hours of RS (**Figure 5**; dark magenta box in **Figure 3**). Terms for upregulated genes requiring 5–6 hours of SD formed clusters enriched in mTOR signaling (e.g., *Pik3ca, Pik3r1*) and cell adhesion (e.g., *Pik3ca, Pik3r1, Crk, Vav3*) (**Figure 5A**). Upregulated genes also belonged to circadian rhythms (e.g., *Clock, Creb1, Npas2*) and stress response (e.g., *Ahsa2, Hsf1, Hspe1*). Clusters of downregulated functional terms included cholesterol biosynthesis (e.g., *Mvd, Msmo1*), ion transport (e.g., *Kcnj4, Pllp*), cell signaling (e.g., *Kcnj4, Gnai2, Prkaca*), and glucose metabolism (e.g., *Ugp2*). Downregulated terms that did not cluster included glycogen biosynthesis (e.g., *Agl, Akt1, Gbe1*). For genes differentially expressed already after 3 hours of SD, clusters of upregulated functional terms included RNA-binding (e.g., *Srsf7, Tra2a, Rbm12b1, Ythdc1*) and stress response (e.g., *Dnajc3, Hspa1b, Hspa5*) (**Figure 5B**). Some upregulated terms, like chromatin regulator (e.g., *Chd9*), did not cluster. Downregulated functional terms were all unclustered and included ubiquitin-like (Ubl) protein conjugation (e.g., *Usp2, Fbxl4, Fbxl21*) and cGMP biosynthesis (e.g., *Npr2, Gucy2e*), which was the term with the highest fold enrichment. Functional enrichment analysis on the union of all genes affected by SD that normalized within 6 hours highlighted stress response (e.g., *Dnajc3, Hspa2b*) and cholesterol biosynthesis (e.g., *Mvd*) as important molecular signatures (**Supplementary Figure S4**).

We then examined functional terms enriched in genes still expressed after 6 hours of RS (**Figure 6**; lavender box in **Figure 3**). Terms for upregulated genes requiring 5–6 hours of SD did not cluster (**Figure 6A**). Unclustered upregulated terms included genes belonging to stress response (e.g., *Ahsa1, Hspa4, Hspb8*) and phagocytosis (e.g., *Elmo1, Elmo2*), which was the term with the highest fold enrichment. Among downregulated genes, there was one cluster that contained genes associated with functional terms involved in cholesterol biosynthesis (e.g., *Nsdhl, Cyb5r3, Mvk, Tm7sf2*). Downregulated functional terms that did not cluster included biosynthesis of cofactors (e.g., *Mmab, Rdh11*) and insulin signaling pathway (e.g., *Flot1, Mapk3, Mknk1*). For DEGs already present after 3 hours of SD, upregulated functional terms clustered into immune response pathways (e.g., *Calr, Cdkn1a, Mapk11*) and stress response (e.g., *Hsp90b1, Hyou1, Hspa8, Dnajb11, Calr*) (**Figure 6B**). Notably, both clusters had high fold enrichments. Upregulated functional terms that did not cluster included MAPK signaling (e.g., *Mapk11, Bdnf)*. The singular downregulated cluster was metabolic pathways (e.g., *Arsb, Galns*). Functional enrichment analysis on the union of all genes affected by SD that remained differentially expressed after 6 hours of RS revealed the MAPK and PI3K/AKT signaling pathways (e.g., *Homer1*) as an additional cluster (**Supplementary Figure S5**).

**Figure 6.**
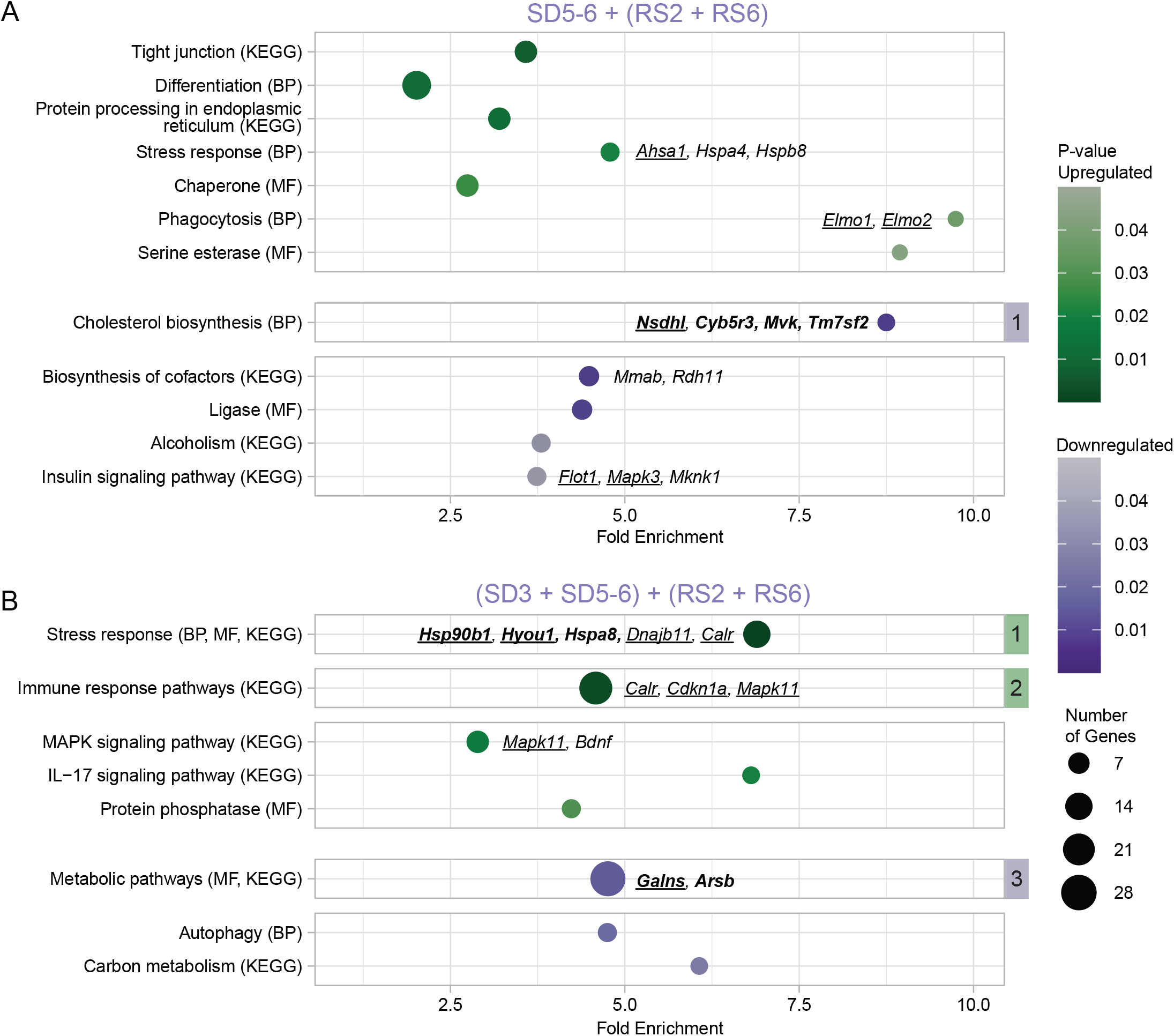
DEGs after SD that are still detected even after 6 hours of RS are involved in stress response, immune response, and metabolic pathways, including cholesterol biosynthesis. Bubble plots of the results of functional enrichment for A) genes differentially expressed at SD5–6, RS2, and RS6 and B) genes differentially expressed at SD3, SD5–6, RS2, and RS6. Enriched (modified Fisher’s Exact p-value < 0.05) functional annotation terms or clusters are displayed vertically and plotted as circles. Clusters are denoted by numbered square boxes to the right of the figure. Functional terms are either Kyoto Encyclopedia of Genes and Genomes (KEGG) pathways, UniProt biological processes (BP), or UniProt molecular functions (MF). Circle size represents the number of genes in a term or a cluster. Upregulated functional terms are shown in green and downregulated terms in purple, with darker shades representing smaller p-values (or the geometric mean of p-values for clusters). Fold enrichment (or the geometric mean of the fold enrichment for clusters) is shown on the x-axis. Hub genes are in bold. Positive control genes are underlined. Representative genes are in regular italics.

### Recovery sleep induces unique gene expression changes that seem to counter the effects of sleep deprivation

Finally, we asked what functional terms were enriched in genes unaffected by SD, but present at a singular RS timepoint (**Figure 7**; dark blue box in **Figure 3**). We first analyzed genes only expressed at RS2, finding that upregulated terms clustered into RNA-binding (e.g., *Srsf1, Rbm8a, Rbm17*) and electron transport (e.g., *Ndufb5, Ndufa8*) (**Figure 7A**). Ribosomal protein (e.g., *Mrpl45, Rps23*) and mRNA transport (e.g., *Rbm8a, Srsf1*) were unclustered upregulated terms. Phosphorylation (e.g., *Mapk12, Map2k3, Map3k9*) was the sole cluster among downregulated terms. Downregulated functional terms that did not cluster included Ubl conjugation (e.g., *Usp5, Usp13, Usp36, Usp49*), Wnt signaling (e.g., *Wnt7b, Ctnnbip1, Sfrp2*), and glycosaminoglycan biosynthesis (e.g., *B4galt2, Chst2, St3gal1*), which was the term with the highest fold enrichment. Subsequently, we focused on genes only expressed at RS6 (**Figure 7B**). Neither upregulated nor downregulated terms clustered. Upregulated functional terms included neurogenesis (e.g., *Adcyap1, Brinp1*), hepatocellular carcinoma (e.g., *Wnt9a*), biological rhythms (e.g., *Calvin3, Timeless*), and tRNA processing (e.g., *Mettl1*).

**Figure 7.**
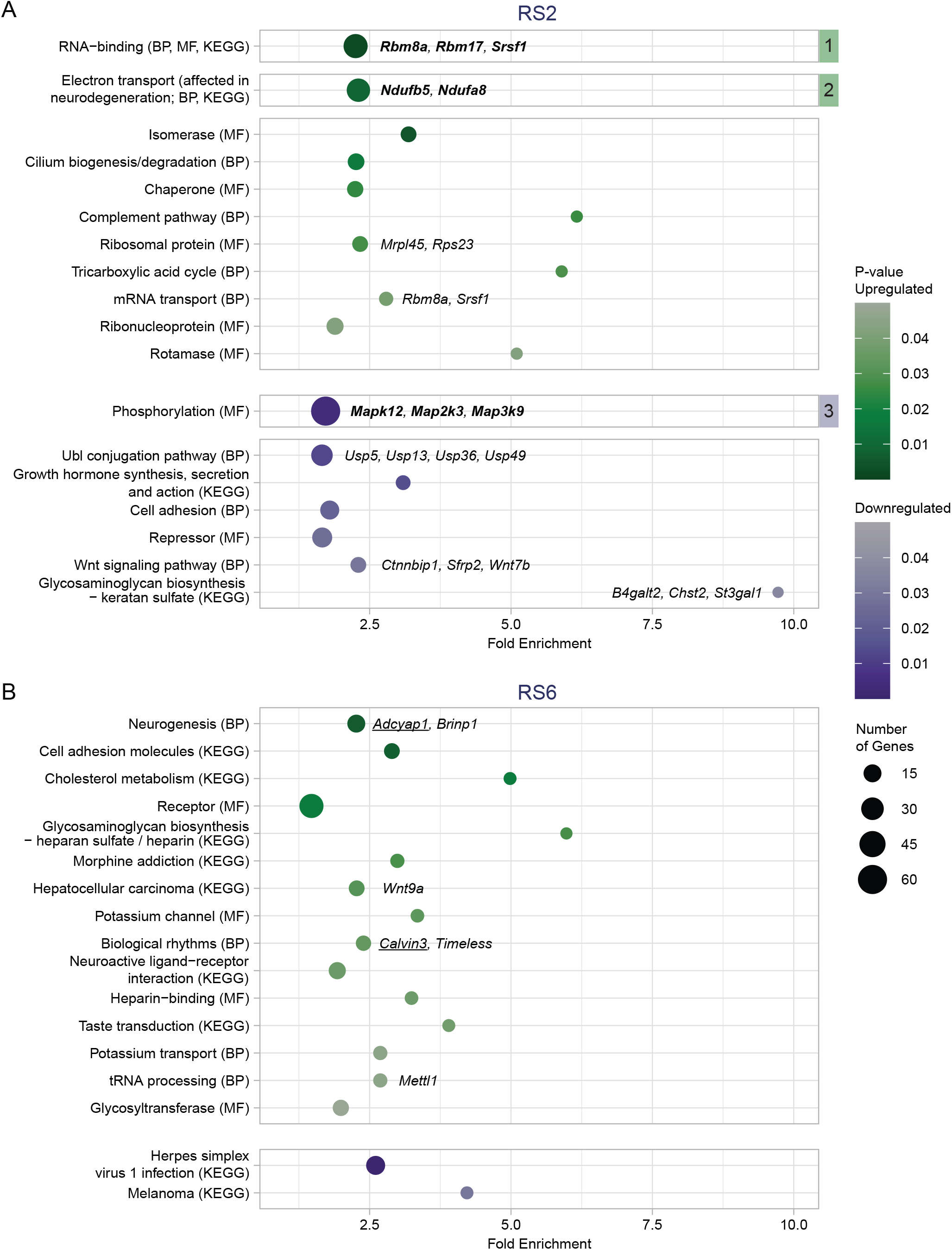
DEGs unique to RS time points are involved in RNA-binding, electron transport, phosphorylation, and ubiquitin-like (Ubl) protein conjugation. Bubble plots of the results of functional enrichment for A) genes differentially expressed only at RS2 and B) genes differentially expressed only at RS6 and SD5–6. Enriched (modified Fisher’s Exact p-value < 0.05) functional annotation terms or clusters are displayed vertically and plotted as circles. Clusters are denoted by numbered square boxes to the right of the figure. Functional terms are either Kyoto Encyclopedia of Genes and Genomes (KEGG) pathways, UniProt biological processes (BP), or UniProt molecular functions (MF). Circle size represents the number of genes in a term or a cluster. Upregulated functional terms are shown in green and downregulated terms in purple, with darker shades representing smaller p-values (or the geometric mean of p-values for clusters). Fold enrichment (or the geometric mean of the fold enrichment for clusters) is shown on the x-axis. Hub genes are in bold. Positive control genes are underlined. Representative genes are in regular italics.

## Discussion

In this study, we examined how genome-wide transcription can be modulated by different amounts of acute SD and subsequent RS in the cortex of adult male WT mice. As we have previously reported, SD has a large and primarily repressive effect on transcription (13). This effect is maximal after 5-6 hours of SD when sleep pressure is also highest (**Figures 2 and 3**). We find that the majority of the DEGs (3,976) are normalized (i.e., no longer detected as differentially expressed) after 2 hours of RS (red box in **Figure 3**). The majority of sleep pressure discharge and most of the adverse effects of SD on hippocampal plasticity are known to revert with 2-3 hours of RS (2, 31, 32). Some recent studies suggest that sleep pressure dissipates even faster, within 60 min of RS (33). Thus, the DEGs after SD that are quickly responsive to RS (**Figure 4**) can provide likely molecular processes involved in sleep homeostasis and SD-dependent impairments. Some of these DEGs require 5-6 hours of SD and include the upregulation of genes involved in oxidative phosphorylation (*mt-Co1, mt-Cytb*), glutathione metabolism (*Gtsm5, Mgst1*), chromatin regulation (*Ctcf, Dnmt3a, Parp1*), and the downregulation of RNA-binding proteins (*Rbm48, Rbm34*) and eukaryotic initiation factors (*Eif4a2, Eif1ax, Eif2ak2*). Some of the quickly responsive DEGs are already observed after only 3 hours of SD. This includes the upregulation of immediate early/MAPK genes (*Nr4a1, Fos, Egr1, Sik1*), and circadian genes (*Per1, Bhlhe40 (Dec1)*), and the downregulation of DNA damage/repair genes (*Rad50* and *Xpa*). In addition, just 2 hours of RS induces gene expression changes not observed after SD, including genes involved in mitochondrial metabolism (*Ndufb5, Ndufa8*) and Wnt signaling (*Wnt7b, Ctnnbip1, Sfrp2*) (**Figure 7**). These genes may underlie additional qualitative differences between baseline and rebound sleep. Finally, some functions and pathways upregulated after SD (see **Figure 4**: phosphorylation, biological rhythms, Ubl conjugation pathway, repressor, and growth hormone metabolism) are found to be downregulated by RS (see **Figure 7**). The same is true for some functions downregulated after SD (see **Figure 4**: RNA-binding, heparin-binding, tRNA processing, and Parkinson’s disease), which are subsequently upregulated by RS. Although these are non-overlapping sets of genes, it suggests that RS may also counter the effects of SD for a subset of pathways and molecular functions. A smaller subset of DEGs after SD (3,047) do not respond readily to RS, taking 6 or more hours of RS to normalize. These genes are involved in cholesterol metabolism (*Mvd, Msmo1*), stress response (*Dnajc3, Hspa1b, Hspa5*), and ion transport (*Kcnj4, Pllp*) (**Figures 5 and 6**). This may reflect non-sleep-specific changes induced by SD, as these DEGs remain after sleep pressure has been discharged. Additionally, we observed pathways and functions altered by SD that contain DEGs at all studied time points, such as insulin signaling and MAPK signaling (**Figures 4-7**), which may simply reflect neuronal activity.

Our findings align with previously published observations and provide further insight into the molecular response to acute SD and subsequent RS. In agreement with our findings, DNA damage as a molecular substrate for sleep pressure has already been described in zebrafish (34). We find rapid downregulation of genes responsible for repairing double-stranded DNA (Rad50, Xpa) after 3 hours of SD, followed at SD5-6 by the upregulation of *Parp1*, which acts as a DNA-damage sensor and is responsible for recruiting DNA repair factors to sites of DNA damage (35). In zebrafish, *Parp1* was shown to increase after SD and induce sleep (34). Thus, our data points to the potential evolutionary conservation of this mechanism as a cellular substrate of sleep pressure. DNA damage is closely linked with redox metabolism (36), as an increase in reactive oxygen species (ROS) can lead to double-stranded DNA breaks. Our study also points to the downregulation of glutathione (a key antioxidant) metabolism as a component of the sleep homeostatic response. In fact, glutathione supplementation can decrease levels of ROS following SD (37) and promote sleep (38). Interestingly, in the same functional cluster as *Parp1 (chromatin regulation)*, we find upregulation of DNA methyltransferase *Dnmt3a* and 3D-chromatin architecture regulator *Ctcf*. This is because *Parp1*, in addition to serving as a DNA-damage sensor, is also responsible for regulating chromatin structure (39). Changes in DNA methylation after SD have been reported in both mice (40) and humans (41), and sleep has been shown to alter 3D nuclear architecture in zebrafish (42). Thus, our results provide a potential link between oxidative stress, DNA damage, DNA methylation and 3D-chromatin conformation as part of the molecular basis of sleep homeostasis. Another subset of genes our study points to as likely to be involved in sleep homeostasis are circadian transcription factors. A link between clock genes and sleep homeostasis has long been hypothesized (43). Our study, however, points to a different subset of clock genes than previously thought: transcriptional repressors *Bhlhe40* (*Dec1*) and *Bhlhe41* (*Dec2*). We find that *Dec1* is quickly upregulated at SD3, increases further at SD5-6, and normalizes after 2 hours of RS. *Dec2* is subsequently downregulated at SD5-6, as expected from being the target of repressor Dec1 (44). *Dec2* mutations are associated with short sleep duration in both humans and mice and lead to reduced repressor activity as well as slightly higher power in the delta range (0-4Hz) of non-rapid eye movement sleep (NREMS) (45). An increase in normalized NREM delta power is considered the gold standard measure of sleep pressure in mammals (2, 46, 47). Thus, perhaps reducing Dec2 expression may be a mechanism by which the increase in NREM delta power after SD is achieved. Last, our results point to a potential role of SD in regulating genes and processes in the frontal cortex that have already been associated with the negative effects of acute SD in the hippocampus, such as insulin/mTOR signaling and the downstream repression of eukaryotic initiation factors (7, 48). This suggests that perhaps the adverse effects of SD in the cortex and hippocampus share common molecular pathways, although no one has ever directly compared the effects of SD on the hippocampus and cortex.

Overall, this study offers new insights into the molecular basis of the homeostatic response to SD as well as SD-dependent impairments by examining combinatorial patterns of gene expression across multiple SD and subsequent RS time points. Genes unique to SD that respond to RS in a similar timeframe as the discharge of sleep pressure include those involved in DNA damage/repair, chromatin regulation, DNA-methylation, redox metabolism and circadian transcription factors. These include genes like *Parp1, Dnmt3a, Ctcf, Dec1 and Dec2* and have a potential role in sleep homeostasis. In contrast, transcriptional changes that occur after SD and are not restored by RS in the timeframe of our study, including the cellular stress response, are more likely to be non-sleep-specific effects of SD. Some limitations curtail the generalizability of our findings, such as the use of only adult male mice. It is thus imperative that future studies include both male and female mice, as well as investigate how dynamics change across different age groups. Last, because this is primarily a hypothesis-generating study, future studies aimed at the validation of the role of genes and pathways identified are warranted.

## Data availability

Sequencing data have been deposited in the Gene Expression Omnibus Database (GEO) under accession number GSE237419. All samples besides those for the RS2 and HC7 time points were previously deposited in GEO under either GSE113754 (SD5, HC5) or GSE140345 (SD3, HC3, SD6, HC6, RS6, and HC12). The code used for the analysis and visualization is available on GitHub at https://github.com/PeixotoLab/SDRS_Transcriptional_Dynamics.

## Supplemental Data

Supplemental Figs. S1–S5 and Supplemental Tables S1–S5: https://10.5281/zenodo.12603038

## Acknowledgments

We would like to acknowledge Christine Mulheim for her contribution to the code used for data analysis.

## Author Contributions

L.P. conceived and designed research; T.W.P., A.I. performed experiments; A.P., S.H. analyzed data; A.P., C.O., interpreted results of experiments; A.P., C.O., K.F. prepared figures; A.P., C.O., L.P. drafted the manuscript; A.P., C.O., K.F., E.M., S.H., K.S., L.P. edited and revised the manuscript; A.P., C.O., K.F., E.M., T.W.P., A.I., S.H., K.S., L.P. approved the final version of the manuscript.

## References

1. Borbély A. The two-process model of sleep regulation: Beginnings and outlook. J Sleep Res 31: e13598, 2022. doi: 10.1111/jsr.13598.

2. Franken P, Chollet D, Tafti M. The homeostatic regulation of sleep need is under genetic control. J Neurosci Off J Soc Neurosci 21: 2610–2621, 2001.

3. Cirelli C, Gutierrez CM, Tononi G. Extensive and divergent effects of sleep and wakefulness on brain gene expression. Neuron 41: 35–43, 2004. doi: 10.1016/s0896-6273(03)00814-6.

4. Mackiewicz M, Shockley KR, Romer MA, Galante RJ, Zimmerman JE, Naidoo N, Baldwin DA, Jensen ST, Churchill GA, Pack AI. Macromolecule biosynthesis: a key function of sleep. Physiol Genomics 31: 441–457, 2007. doi: 10.1152/physiolgenomics.00275.2006.

5. Maret S, Dorsaz S, Gurcel L, Pradervand S, Petit B, Pfister C, Hagenbuchle O, O’Hara BF, Franken P, Tafti M. Homer1a is a core brain molecular correlate of sleep loss. Proc Natl Acad Sci U S A 104: 20090–20095, 2007. doi: 10.1073/pnas.0710131104.

6. Thompson CL, Wisor JP, Lee C-K, Pathak SD, Gerashchenko D, Smith KA, Fischer SR, Kuan CL, Sunkin SM, Ng LL, Lau C, Hawrylycz M, Jones AR, Kilduff TS, Lein ES. Molecular and anatomical signatures of sleep deprivation in the mouse brain. Front Neurosci 4: 165, 2010. doi: 10.3389/fnins.2010.00165.

7. Vecsey CG, Peixoto L, Choi JHK, Wimmer M, Jaganath D, Hernandez PJ, Blackwell J, Meda K, Park AJ, Hannenhalli S, Abel T. Genomic analysis of sleep deprivation reveals translational regulation in the hippocampus. Physiol Genomics 44: 981–991, 2012. doi: 10.1152/physiolgenomics.00084.2012.

8. Gerstner JR, Koberstein JN, Watson AJ, Zapero N, Risso D, Speed TP, Frank MG, Peixoto L. Removal of unwanted variation reveals novel patterns of gene expression linked to sleep homeostasis in murine cortex. BMC Genomics 17: 727, 2016. doi: 10.1186/s12864-016-3065-8.

9. Noya SB, Colameo D, Brüning F, Spinnler A, Mircsof D, Opitz L, Mann M, Tyagarajan SK, Robles MS, Brown SA. The forebrain synaptic transcriptome is organized by clocks but its proteome is driven by sleep. Science 366: eaav2642, 2019. doi: 10.1126/science.aav2642.

10. Gaine ME, Bahl E, Chatterjee S, Michaelson JJ, Abel T, Lyons LC. Altered hippocampal transcriptome dynamics following sleep deprivation. Mol Brain 14: 125, 2021. doi: 10.1186/s13041-021-00835-1.

11. Muheim CM, Ford K, Medina E, Singletary K, Peixoto L, Frank MG. Ontogenesis of the molecular response to sleep loss. Neurobiol Sleep Circadian Rhythms 14: 100092, 2023. doi: 10.1016/j.nbscr.2023.100092.

12. Vanrobaeys Y, Mukherjee U, Langmack L, Beyer SE, Bahl E, Lin L-C, Michaelson JJ, Abel T, Chatterjee S. Mapping the spatial transcriptomic signature of the hippocampus during memory consolidation. Nat Commun 14: 6100, 2023. doi: 10.1038/s41467-023-41715-7.

13. Ford K, Zuin E, Righelli D, Medina E, Schoch H, Singletary K, Muheim C, Frank MG, Hicks SC, Risso D, Peixoto L. A Global Transcriptional Atlas of the Effect of Sleep Deprivation in the Mouse Frontal Cortex. bioRxiv: 2023.11.28.569011, 2023.

14. Hor CN, Yeung J, Jan M, Emmenegger Y, Hubbard J, Xenarios I, Naef F, Franken P. Sleep-wake-driven and circadian contributions to daily rhythms in gene expression and chromatin accessibility in the murine cortex. Proc Natl Acad Sci U S A 116: 25773–25783, 2019. doi: 10.1073/pnas.1910590116.

15. Xu Z-Q, Gao C-Y, Fang C-Q, Zhou H-D, Jiang X-J. The mechanism and characterization of learning and memory impairment in sleep-deprived mice. Cell Biochem Biophys 58: 137–140, 2010. doi: 10.1007/s12013-010-9098-8.

16. Hagewoud R, Havekes R, Novati A, Keijser JN, Van der Zee EA, Meerlo P. Sleep deprivation impairs spatial working memory and reduces hippocampal AMPA receptor phosphorylation. J Sleep Res 19: 280–288, 2010. doi: 10.1111/j.1365-2869.2009.00799.x.

17. Ingiosi AM, Schoch H, Wintler T, Singletary KG, Righelli D, Roser LG, Medina E, Risso D, Frank MG, Peixoto L. Shank3 modulates sleep and expression of circadian transcription factors. eLife 8: e42819, 2019. doi: 10.7554/eLife.42819.

18. Patro R, Duggal G, Love MI, Irizarry RA, Kingsford C. Salmon provides fast and bias-aware quantification of transcript expression. Nat Methods 14: 417–419, 2017. doi: 10.1038/nmeth.4197.

19. Love MI, Soneson C, Hickey PF, Johnson LK, Pierce NT, Shepherd L, Morgan M, Patro R. Tximeta: Reference sequence checksums for provenance identification in RNA-seq. PLoS Comput Biol 16: e1007664, 2020. doi: 10.1371/journal.pcbi.1007664.

20. Gentleman RC, Carey VJ, Bates DM, Bolstad B, Dettling M, Dudoit S, Ellis B, Gautier L, Ge Y, Gentry J, Hornik K, Hothorn T, Huber W, Iacus S, Irizarry R, Leisch F, Li C, Maechler M, Rossini AJ, Sawitzki G, Smith C, Smyth G, Tierney L, Yang JYH, Zhang J. Bioconductor: open software development for computational biology and bioinformatics. Genome Biol 5: R80, 2004. doi: 10.1186/gb-2004-5-10-r80.

21. Huber W, Carey VJ, Gentleman R, Anders S, Carlson M, Carvalho BS, Bravo HC, Davis S, Gatto L, Girke T, Gottardo R, Hahne F, Hansen KD, Irizarry RA, Lawrence M, Love MI, MacDonald J, Obenchain V, Oleś AK, Pagès H, Reyes A, Shannon P, Smyth GK, Tenenbaum D, Waldron L, Morgan M. Orchestrating high-throughput genomic analysis with Bioconductor. Nat Methods 12: 115–121, 2015. doi: 10.1038/nmeth.3252.

22. Risso D, Schwartz K, Sherlock G, Dudoit S. GC-Content Normalization for RNA-Seq Data. BMC Bioinformatics 12: 480, 2011. doi: 10.1186/1471-2105-12-480.

23. Risso D, Ngai J, Speed TP, Dudoit S. Normalization of RNA-seq data using factor analysis of control genes or samples. Nat Biotechnol 32: 896–902, 2014. doi: 10.1038/nbt.2931.

24. Robinson MD, McCarthy DJ, Smyth GK. edgeR: a Bioconductor package for differential expression analysis of digital gene expression data. Bioinformatics 26: 139–140, 2010. doi: 10.1093/bioinformatics/btp616.

25. Martin FJ, Amode MR, Aneja A, Austine-Orimoloye O, Azov AG, Barnes I, Becker A, Bennett R, Berry A, Bhai J, Bhurji SK, Bignell A, Boddu S, Branco Lins PR, Brooks L, Ramaraju SB, Charkhchi M, Cockburn A, Da Rin Fiorretto L, Davidson C, Dodiya K, Donaldson S, El Houdaigui B, El Naboulsi T, Fatima R, Giron CG, Genez T, Ghattaoraya GS, Martinez JG, Guijarro C, Hardy M, Hollis Z, Hourlier T, Hunt T, Kay M, Kaykala V, Le T, Lemos D, Marques-Coelho D, Marugán JC, Merino GA, Mirabueno LP, Mushtaq A, Hossain SN, Ogeh DN, Sakthivel MP, Parker A, Perry M, Piližota I, Prosovetskaia I, Pérez-Silva JG, Salam AIA, Saraiva-Agostinho N, Schuilenburg H, Sheppard D, Sinha S, Sipos B, Stark W, Steed E, Sukumaran R, Sumathipala D, Suner M-M, Surapaneni L, Sutinen K, Szpak M, Tricomi FF, Urbina-Gómez D, Veidenberg A, Walsh TA, Walts B, Wass E, Willhoft N, Allen J, Alvarez-Jarreta J, Chakiachvili M, Flint B, Giorgetti S, Haggerty L, Ilsley GR, Loveland JE, Moore B, Mudge JM, Tate J, Thybert D, Trevanion SJ, Winterbottom A, Frankish A, Hunt SE, Ruffier M, Cunningham F, Dyer S, Finn RD, Howe KL, Harrison PW, Yates AD, Flicek P. Ensembl 2023. Nucleic Acids Res 51: D933–D941, 2022. doi: 10.1093/nar/gkac958.

26. Conway JR, Lex A, Gehlenborg N. UpSetR: an R package for the visualization of intersecting sets and their properties. Bioinformatics 33: 2938–2940, 2017. doi: 10.1093/bioinformatics/btx364.

27. Huang DW, Sherman BT, Lempicki RA. Systematic and integrative analysis of large gene lists using DAVID bioinformatics resources. Nat Protoc 4: 44–57, 2009. doi: 10.1038/nprot.2008.211.

28. Sherman BT, Hao M, Qiu J, Jiao X, Baseler MW, Lane HC, Imamichi T, Chang W. DAVID: a web server for functional enrichment analysis and functional annotation of gene lists (2021 update). Nucleic Acids Res 50: W216–W221, 2022. doi: 10.1093/nar/gkac194.

29. Peixoto L, Risso D, Poplawski SG, Wimmer ME, Speed TP, Wood MA, Abel T. How data analysis affects power, reproducibility and biological insight of RNA-seq studies in complex datasets. Nucleic Acids Res 43: 7664–7674, 2015. doi: 10.1093/nar/gkv736.

30. Hor CN, Yeung J, Jan M, Emmenegger Y, Hubbard J, Xenarios I, Naef F, Franken P. Sleep-wake-driven and circadian contributions to daily rhythms in gene expression and chromatin accessibility in the murine cortex. Proc Natl Acad Sci U S A 116: 25773–25783, 2019. doi: 10.1073/pnas.1910590116.

31. Havekes R, Park AJ, Tudor JC, Luczak VG, Hansen RT, Ferri SL, Bruinenberg VM, Poplawski SG, Day JP, Aton SJ, Radwańska K, Meerlo P, Houslay MD, Baillie GS, Abel T. Sleep deprivation causes memory deficits by negatively impacting neuronal connectivity in hippocampal area CA1. eLife 5: e13424, 2016. doi: 10.7554/eLife.13424.

32. Vecsey CG, Baillie GS, Jaganath D, Havekes R, Daniels A, Wimmer M, Huang T, Brown KM, Li X-Y, Descalzi G, Kim SS, Chen T, Shang Y-Z, Zhuo M, Houslay MD, Abel T. Sleep deprivation impairs cAMP signaling in the hippocampus. Nature 461: 1122–1125, 2009. doi: 10.1038/nature08488.

33. Hubbard J, Gent TC, Hoekstra MMB, Emmenegger Y, Mongrain V, Landolt H-P, Adamantidis AR, Franken P. Rapid fast-delta decay following prolonged wakefulness marks a phase of wake-inertia in NREM sleep. Nat Commun 11: 3130, 2020. doi: 10.1038/s41467-020-16915-0.

34. Zada D, Sela Y, Matosevich N, Monsonego A, Lerer-Goldshtein T, Nir Y, Appelbaum L. Parp1 promotes sleep, which enhances DNA repair in neurons. Mol Cell 81: 4979–4993.e7, 2021. doi: 10.1016/j.molcel.2021.10.026.

35. Gibson BA, Kraus WL. New insights into the molecular and cellular functions of poly(ADP-ribose) and PARPs. Nat Rev Mol Cell Biol 13: 411–424, 2012. doi: 10.1038/nrm3376.

36. Shadfar S, Parakh S, Jamali MS, Atkin JD. Redox dysregulation as a driver for DNA damage and its relationship to neurodegenerative diseases. Transl Neurodegener 12: 18, 2023. doi: 10.1186/s40035-023-00350-4.

37. Villafuerte G, Miguel-Puga A, Murillo Rodríguez E, Machado S, Manjarrez E, Arias-Carrión O. Sleep Deprivation and Oxidative Stress in Animal Models: A Systematic Review. Oxid Med Cell Longev 2015: 234952, 2015. doi: 10.1155/2015/234952.

38. Kimura M, Kapás L, Krueger JM. Oxidized glutathione promotes sleep in rabbits. Brain Res Bull 45: 545–548, 1998. doi: 10.1016/s0361-9230(97)00441-3.

39. Kraus WL. Transcriptional control by PARP-1: chromatin modulation, enhancer-binding, coregulation, and insulation. Curr Opin Cell Biol 20: 294–302, 2008. doi: 10.1016/j.ceb.2008.03.006.

40. Massart R, Freyburger M, Suderman M, Paquet J, El Helou J, Belanger-Nelson E, Rachalski A, Koumar OC, Carrier J, Szyf M, Mongrain V. The genome-wide landscape of DNA methylation and hydroxymethylation in response to sleep deprivation impacts on synaptic plasticity genes. Transl Psychiatry 4: e347–e347, 2014. doi: 10.1038/tp.2013.120.

41. Trivedi MS, Holger D, Bui AT, Craddock TJA, Tartar JL. Short-term sleep deprivation leads to decreased systemic redox metabolites and altered epigenetic status. PLoS ONE 12: e0181978, 2017. doi: 10.1371/journal.pone.0181978.

42. Zada D, Bronshtein I, Lerer-Goldshtein T, Garini Y, Appelbaum L. Sleep increases chromosome dynamics to enable reduction of accumulating DNA damage in single neurons. Nat Commun 10: 895, 2019. doi: 10.1038/s41467-019-08806-w.

43. Franken P, Dijk D-J. Circadian clock genes and sleep homeostasis. Eur J Neurosci 29: 1820–1829, 2009. doi: 10.1111/j.1460-9568.2009.06723.x.

44. Li Y, Xie M, Song X, Gragen S, Sachdeva K, Wan Y, Yan B. DEC1 Negatively Regulates the Expression of DEC2 through Binding to the E-box in the Proximal Promoter*. J Biol Chem 278: 16899–16907, 2003. doi: 10.1074/jbc.M300596200.

45. He Y, Jones CR, Fujiki N, Xu Y, Guo B, Holder JL, Rossner MJ, Nishino S, Fu Y-H. The Transcriptional Repressor DEC2 Regulates Sleep Length in Mammals. Science 325: 866–870, 2009. doi: 10.1126/science.1174443.

46. Achermann P, Borbély AA. Mathematical models of sleep regulation. Front Biosci J Virtual Libr 8: s683–693, 2003. doi: 10.2741/1064.

47. Borb AA, Achermann P. Sleep Homeostasis and Models of Sleep Regulation. J Biol Rhythms 14: 559–570, 1999. doi: 10.1177/074873099129000894.

48. Tudor JC, Davis EJ, Peixoto L, Wimmer ME, van Tilborg E, Park AJ, Poplawski SG, Chung CW, Havekes R, Huang J, Gatti E, Pierre P, Abel T. Sleep deprivation impairs memory by attenuating mTORC1-dependent protein synthesis. Sci Signal 9: ra41, 2016. doi: 10.1126/scisignal.aad4949.

