## Supplemental Figures S1-S5 for "Transcriptional dynamics of sleep deprivation and subsequent recovery sleep in the male mouse cortex"

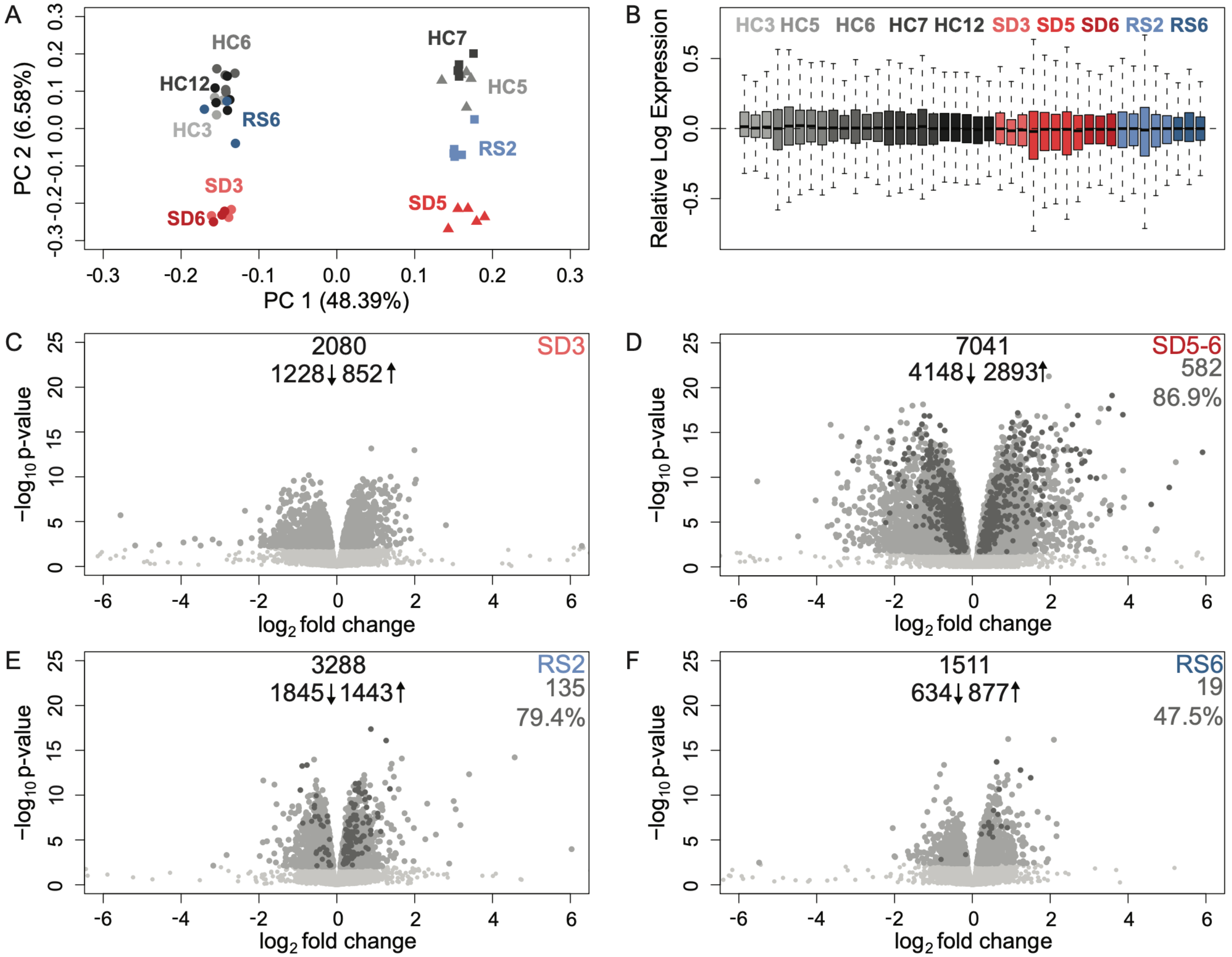


**Supplemental Figure S1. Traditional normalization methods result in fewer differentially expressed genes and positive controls recovered.** A) Principal component analysis following upper-quartile (UQ) normalization. Circles represent samples from (PMID: 31776259), triangles represent samples from (30973326), and squares represent new samples from our group. Home cage (HC) samples are in shades of gray, sleep deprivation (SD) samples are in shades of red, and recovery sleep (RS) samples are in shades of blue. The lab effect (PC1, 48.39%) accounts for nearly half of the variability in the dataset, whereas the amount of sleep only explained 6.58% (PC2). B) Relative log expression for each sample and condition. Color code as in A. C–F) Volcano plots following differential analysis on UQ normalized counts. C) SD3 vs HC3, D) SD5–6 vs HC5–6, E) RS2 vs HC7, F) RS6 vs HC12. Expressed genes are in light gray, differentially expressed genes (DEGs) are in gray, and positive controls are in dark gray. Numbers of downregulated DEGs, upregulated DEGs, and total DEGs are reported in the top center for each comparison. The number and percentage of positive controls recovered are reported in the top right for each comparison.

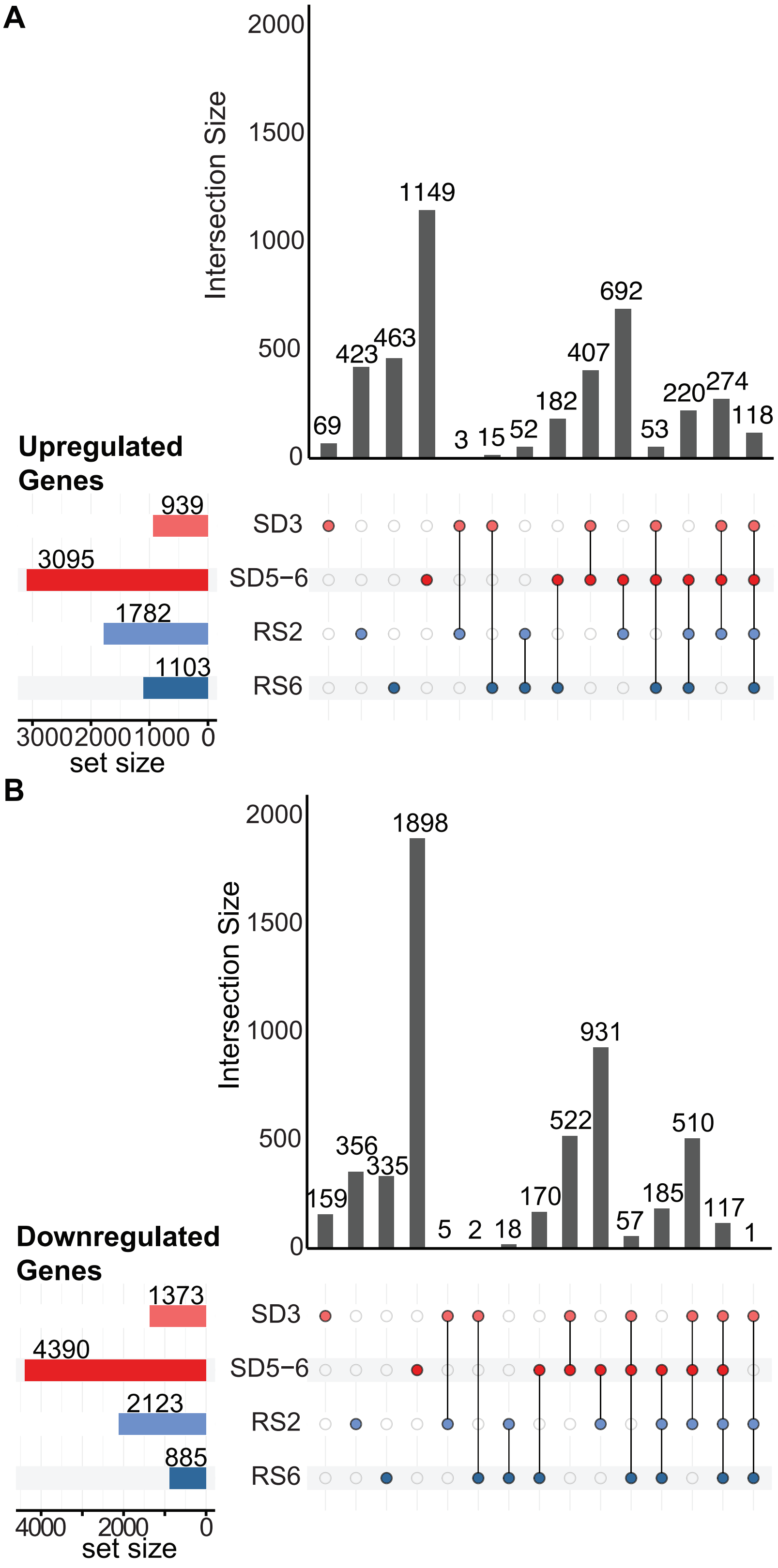


**Supplemental Figure S2. Most differentially expressed genes follow biologically meaningful patterns of expression across time points.** UpSet plots of all non-empty intersections across the SD3, SD5–6, RS2, and RS6 time points. Lists of differentially expressed genes (DEGs) were intersected separately for A) upregulated genes and B) downregulated genes. The total number of DEGs per time point is shown on the set size rows. The vertical bars represent the number of DEGs belonging to the subset depicted by the colored dots in the intersection matrix.

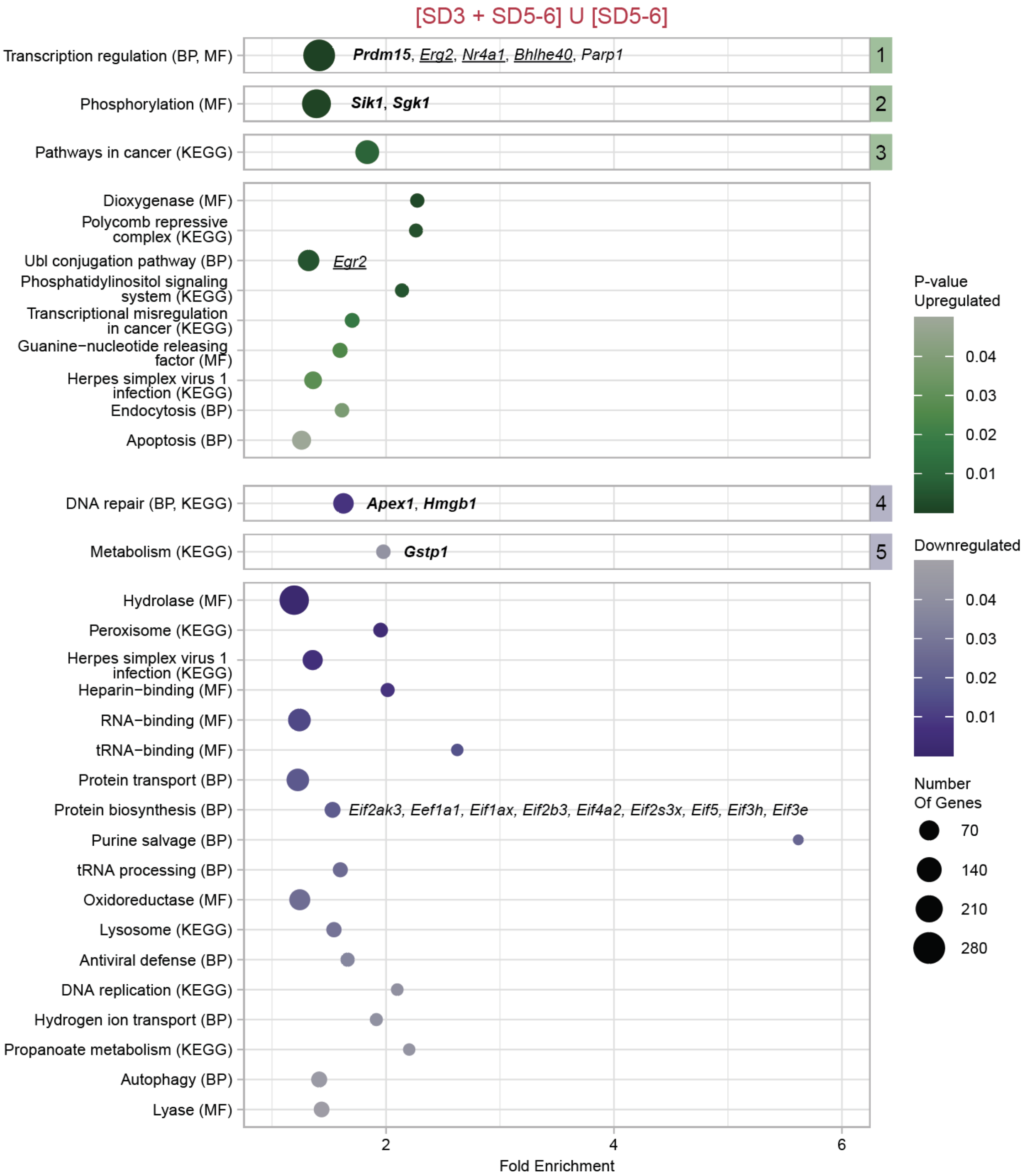


**Supplemental Figure S3. DEGs after SD that are no longer detected after 2 hours of RS are collectively involved in metabolism and protein biosynthesis.** Bubble plot of the results of functional enrichment for genes recovered rapidly after being differentially expressed at either SD3 and SD5–6 or SD5–6 only. Enriched (modified Fisher's Exact p-value < 0.05) functional annotation terms or clusters are displayed vertically and plotted as circles. Clusters are denoted by numbered square boxes to the right of the figure. Functional terms are either Kyoto Encyclopedia of Genes and Genomes (KEGG) pathways, UniProt biological processes (BP), or UniProt molecular functions (MF). Circle size represents the number of genes in a term or a cluster. Upregulated functional terms are shown in green and downregulated terms in purple, with darker shades representing smaller p-values (or the geometric mean of p-values for clusters). Fold enrichment (or the geometric mean of the fold enrichment for clusters) is shown in the x-axis. Hub genes are in bold. Positive control genes are underlined. Representative genes are in regular italics.

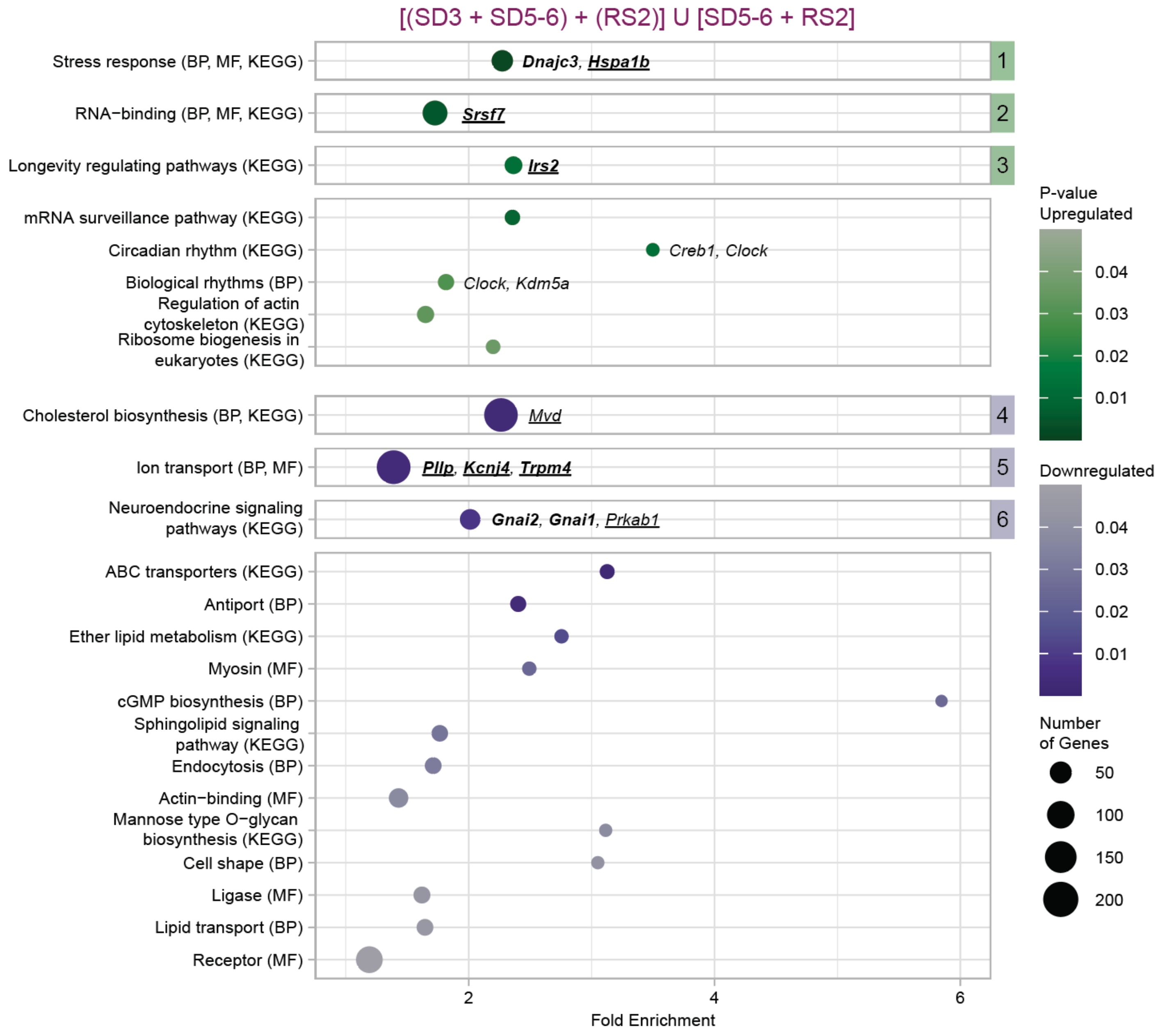


**Supplemental Figure S4. DEGs after SD that are no longer detected after 2–6 hours of RS are collectively involved in stress response and cholesterol biosynthesis.** Bubble plot of the results of functional enrichment. Enriched (modified Fisher's Exact p-value < 0.05) functional annotation terms or clusters are displayed vertically and plotted as circles. Clusters are denoted by numbered square boxes to the right of the figure. Functional terms are either Kyoto Encyclopedia of Genes and Genomes (KEGG) pathways, UniProt biological processes (BP), or UniProt molecular functions (MF). Circle size represents the number of genes in a term or a cluster. Upregulated functional terms are shown in green and downregulated terms in purple, with darker shades representing smaller p-values (or the geometric mean of p-values for clusters). Fold enrichment (or the geometric mean of the fold enrichment for clusters) is shown in the x-axis. Hub genes are in bold. Positive control genes are underlined. Representative genes are in regular italics.

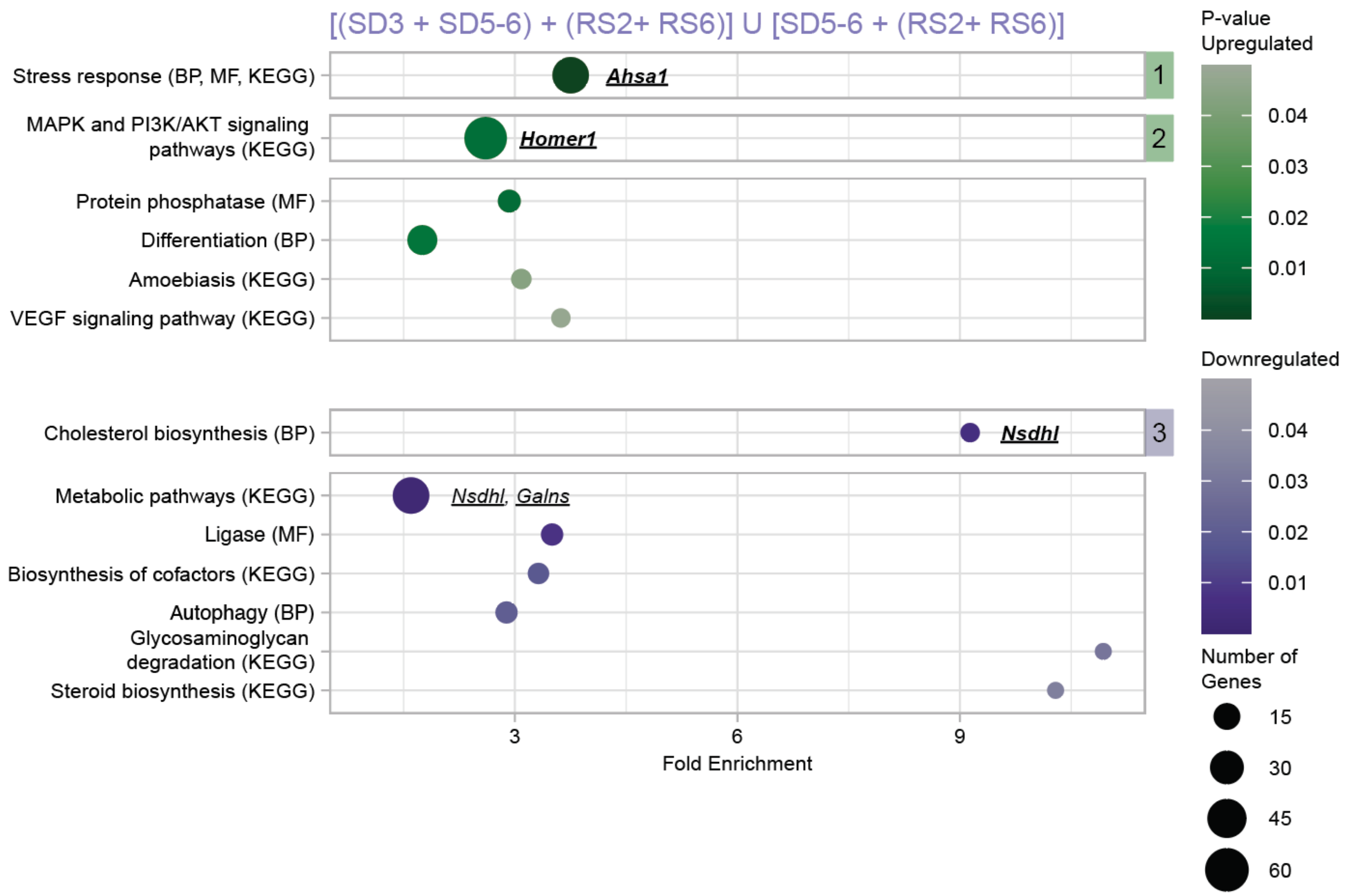


**Supplemental Figure S5. DEGs after SD that are still detected even after 6 hours of RS are collectively involved in metabolic pathways and the MAPK and PI3K/AKT signaling pathways.** Bubble plot of the results of functional enrichment. Enriched (modified Fisher's Exact p-value < 0.05) functional annotation terms or clusters are displayed vertically and plotted as circles. Clusters are denoted by numbered square boxes to the right of the figure. Functional terms are either Kyoto Encyclopedia of Genes and Genomes (KEGG) pathways, UniProt biological processes (BP), or UniProt molecular functions (MF). Circle size represents the number of genes in a term or a cluster. Upregulated functional terms are shown in green and downregulated terms in purple, with darker shades representing smaller p-values (or the geometric mean of p-values for clusters). Fold enrichment (or the geometric mean of the fold enrichment for clusters) is shown in the x-axis. Hub genes are in bold. Positive control genes are underlined. Representative genes are in regular italics.
